# Microtubule poleward flux in human cells is driven by the coordinated action of four kinesins

**DOI:** 10.1101/2020.06.16.155259

**Authors:** Yulia Steblyanko, Girish Rajendraprasad, Mariana Osswald, Susana Eibes, Stephan Geley, António J. Pereira, Helder Maiato, Marin Barisic

**Affiliations:** Danish Cancer Society Research Center (DCRC), Strandboulevarden 49, 2100 Copenhagen, Denmark; i3S - Instituto de Investigação e Inovação em Saúde, Universidade do Porto, Rua Alfredo Allen 208, 4200-135 Porto, Portugal; IBMC - Instituto de Biologia Molecular e Celular, Universidade do Porto, Rua Alfredo Allen 208, 4200-135 Porto, Portugal; Institute of Pathophysiology, Biocenter, Medical University of Innsbruck, Innrain 80, 6020 Innsbruck, Austria; Experimental Biology Unit, Department of Biomedicine, Faculdade de Medicina, Universidade do Porto, Alameda Prof. Hernâni Monteiro, 4200-319 Porto, Portugal; Department of Cellular and Molecular Medicine, Faculty of Health Sciences, University of Copenhagen, Blegdamsvej 3B, 2200 Copenhagen, Denmark

**Keywords:** Kinesins, Kinetochore, Microtubules, Mitosis, Mitotic spindle

## Abstract

Mitotic spindle microtubules (MTs) undergo continuous poleward flux, whose driving force and function in humans remain unclear. Here, we combined loss-of-function screenings with analysis of MT dynamics in human cells to investigate the molecular mechanisms underlying MT-flux. We report that kinesin-7/CENP-E at kinetochores (KTs) is the predominant driver of MT-flux in early prometaphase, while kinesin-4/KIF4A on chromosome arms facilitates MT-flux during late prometaphase and metaphase. We show that both of these activities work in coordination with MT-crosslinking motors kinesin-5/EG5 and kinesin-12/KIF15. Our data further indicate that MT-flux driving force is transmitted from non-KT MTs to KT-MTs via MT-coupling by HSET and NuMA. Moreover, we found that MT-flux rate correlates with spindle size and this correlation depends on the establishment of stable end-on KT-MT attachments. Strikingly, we revealed that flux is required to counteract the kinesin 13/MCAK-dependent MT-depolymerization to regulate spindle length. Thus, our study demonstrates that MT-flux in human cells is driven by the coordinated action of four kinesins, and is required to regulate mitotic spindle size in response to MCAK-mediated MT-depolymerizing activity at KTs.

## Introduction

Microtubule (MT) poleward flux is an evolutionarily conserved process in metazoan spindles, defined as a continuous motion of MTs towards the spindle poles (Bajer & Molè-Bajer, 1972, Forer, 1965, Hamaguchi, Toriyama et al., 1987, Hiramoto & Izutsu, 1977, Mitchison, 1989). Although its function remains unclear, MT-flux has been proposed to play a role in various cellular processes, including regulation of spindle length (Fu, Bian et al., 2015, Gaetz & Kapoor, 2004, Renda, Pellacani et al., 2017, Rogers, Rogers et al., 2004). However, whereas reduced MT-flux led to spindle elongation in *Drosophila* embryos (Rogers et al., 2004) and *Xenopus* egg extracts (Gaetz & Kapoor, 2004), its reduction in human cells either had no effect on spindle length (Ganem, Upton et al., 2005, Jiang, Rezabkova et al., 2017), or it resulted in shorter spindles (Fu et al., 2015, Maffini, Maia et al., 2009), leaving the role of MT-flux in regulation of spindle length ill-defined. Other proposed cellular functions for MT-flux include regulation of kinetochore (KT) activity (Maddox, Straight et al., 2003) and chromosome movements (Ganem et al., 2005, Rogers et al., 2004), correction of erroneous KT-MT attachments (Ganem et al., 2005), and equalization of the forces at KTs prior to their segregation (Matos, Pereira et al., 2009).

In addition to its function, the molecular mechanism underlying MT-flux remains controversial as well. Two main models have been proposed to drive spindle MT-flux. The first model envisions MT-flux to be driven by kinesin-13-induced depolymerization of MT minus-ends at the spindle poles (Ganem et al., 2005, Rogers et al., 2004), coupled to CLASPs-mediated MT polymerization at KTs (Maiato, Khodjakov et al., 2005). However, several lines of evidence have challenged this model. In particular, fluorescence speckle microscopy in newt lung cells showed no flux in astral MTs (Waterman-Storer, Desai et al., 1998), which originate at the poles and extend towards the cell cortex. Moreover, laser microsurgery experiments on KT-MTs (also called k-fibres) revealed normal MT-flux despite stable MT minus-ends detached from the spindle poles (Maiato, Rieder et al., 2004, Matos et al., 2009). Finally, MTs continued to flux at unchanged rates even when MT depolymerization at spindle poles was inhibited by controlled mechanical compression applied to metaphase mitotic spindles (Dumont & Mitchison, 2009). Together, these experiments strongly suggest that MT minus-end depolymerization is a reaction to flux in order to regulate spindle length, rather than the main driving force.

The second model proposes that MT depolymerization and polymerization are a response to kinesin-5-mediated sliding of anti-parallel interpolar MTs (Brust-Mascher & Scholey, 2002, Matos et al., 2009, Miyamoto, Perlman et al., 2004, Pereira & Maiato, 2012), which could then be translated from interpolar-to KT-MTs via MT cross-linking molecules (Shimamoto, Maeda et al., 2011, Vladimirou, McHedlishvili et al., 2013). However, this mechanism was challenged by experiments showing that inhibition of kinesin-5 (EG5 in humans) only slightly reduced MT-flux rates both in bipolar and monopolar spindles (Cameron, Yang et al., 2006). Moreover, the ability of monopolar spindles to flux (Cameron et al., 2006) also eliminated the possibility that antiparallel interpolar MTs play an essential role in this cellular process in mammalian cells.

Thus, although MT-flux has been studied for three decades, the molecular mechanisms underlying spindle MT-flux, as well as its cellular function, remain to be elucidated. In this study, we demonstrate that MT-flux is sequentially driven from prometaphase to metaphase by CENP-E/kinesin-7 at KTs and KIF4A/kinesin-4 on chromosome arms, respectively, and that both of these kinesins work in cooperation with the tetrameric MT-crosslinking motors EG5 and KIF15. These activities are coordinated with the action of the MT-crosslinking molecules HSET and NuMA, which couple non-KT-MTs to KT-MTs, thereby ensuring a uniform distribution of MT-flux-dependent forces across the mitotic spindle. In addition, we shed new light on the cellular function of MT-flux in human cells in the regulation of mitotic spindle size upon the establishment of stable end-on attachments by counteracting MCAK-mediated MT-depolymerizing activity (Desai, Verma et al., 1999) at KTs.

## Results

### KIF4A strongly contributes to MT-flux

In order to gain insight into the molecular mechanisms underlying MT-flux in human somatic cells, we combined RNAi (**Fig EV1A**) and chemical inhibitors with photoactivation-based spinning disk confocal live-cell imaging of late prometaphase/metaphase bipolar spindles and S-trityl-L-cysteine (STLC)-treated monopolar spindles in human osteosarcoma U2OS cells stably expressing photoactivatable (PA)-GFP tubulin (**Fig 1A-C, Movie EV1 and Movie EV2**). We tested established contributors to MT-flux (kinesin-13/KIF2A, CLASP1+2 and kinesin-5/EG5), as well as several additional potential candidates (kinesin-12/KIF15, kinesin-7/CENP-E, kinesin-10/hKID and kinesin-4/KIF4A). KIF2A was investigated because of its ability to depolymerize MT minus-ends at the spindle poles and because of its suggested role in MT-flux both in humans and flies (Ems-McClung & Walczak, 2010), whereas CLASPs contribute to MT polymerization at the KT (Maffini et al., 2009, Maiato et al., 2005). hKID and KIF4A are chromokinesins, plus-end directed motor proteins localized on chromosome arms that generate chromosomal polar ejection forces (PEFs), promoting proper positioning of chromosome arms and accurate chromosome alignment and segregation (Antonio, Ferby et al., 2000, Barisic, Aguiar et al., 2014, Brouhard & Hunt, 2005, Dong, Zhu et al., 2018, Funabiki & Murray, 2000, Levesque & Compton, 2001, Mazumdar, Sundareshan et al., 2004, Rieder, Davison et al., 1986, Rieder & Salmon, 1994, Tipton, Wren et al., 2017, Vernos, Raats et al., 1995, Wandke, Barisic et al., 2012). KIF4A also interacts with PRC1 at interpolar-MTs, facilitating accurate cytokinesis (Zhu & Jiang, 2005). CENP-E is a KT- and MT-localized plus-end directed motor protein required for congression of peripheral chromosomes and proper chromosome segregation (Barisic et al., 2014, Kapoor, Lampson et al., 2006, Schaar, Chan et al., 1997, Wood, Sakowicz et al., 1997, Yen, Compton et al., 1991). KIF15 is another plus-end directed motor with a tetrameric conformation and the ability to slide anti-parallel MTs in a way similar to EG5 (Drechsler, McHugh et al., 2014, Tanenbaum, Macurek et al., 2009). The analysis of sum-projected kymographs (**Fig 1D**) excluded EG5, KIF15 and hKID as strong individual contributors to MT-flux (**Fig 1E and F**). However, depletion of KIF4A revealed a strong contribution to MT-flux, not only in bipolar spindles (0.32 +/− 0.14 μm/min, compared to 0.61 +/− 0.22 μm/min in controls), as we have previously reported (Wandke et al., 2012), but also in monopolar spindles (0.33 +/− 0.11 μm/min, compared to 0.5 +/− 0.14 μm/min in controls). Intriguingly, while inhibition of CENP-E did not significantly affect MT-flux in bipolar spindles (0.52 +/− 0.21 μm/min in this study, and as shown before (Logarinho, Maffini et al., 2012)), we detected a significant reduction of flux in monopolar spindles (0.34 +/− 0.07 μm/min) (**Fig 1E and F**). As expected, depletion of KIF2A and CLASPs led to a strong reduction in flux rates in bipolar (0.17 +/− 0.12 μm/min and 0.27 +/− 0.18 μm/min, respectively) and monopolar spindles (0.17 +/− 0.09 μm/min and 0.28 +/− 0.07 μm/min, respectively) (**Fig 1E and F**), confirming their importance in MT-minus-end depolymerization and plus-end polymerization, respectively.

**Figure 1.**
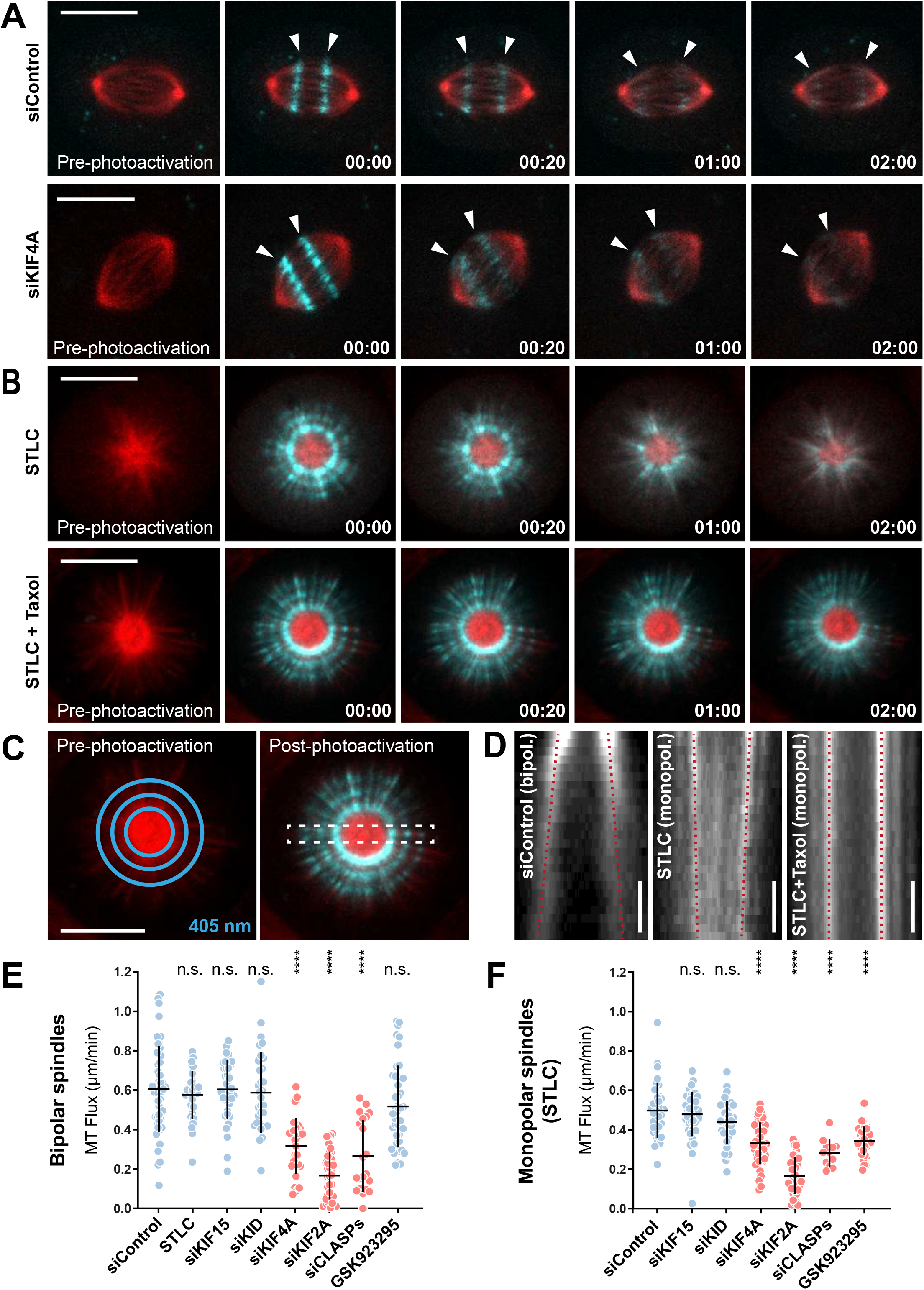
KIF4A strongly contributes to MT-flux. **A.** Representative spinning disk confocal live-cell imaging time series of U2OS cells stably co-expressing PA-GFP-α-tubulin (cyan) and mCherry-α-tubulin (red), treated with control and KIF4A siRNAs. White arrowheads highlight poleward motion of the photoactivated regions due to MT-flux. Scale bars, 10 μm. Time, min:sec. **B.** Representative spinning disk confocal live-cell imaging time series images of S-trityl-L-cysteine (STLC)-treated U2OS cells stably co-expressing PA-GFP-α-tubulin (cyan) and mCherry-α-tubulin (red). Note that MT-flux is abrogated in presence of taxol (lower panel). Scale bars, 10 μm. Time, min:sec. **C.** Illustration of 405nm laser-photoactivated regions in monopolar spindles (blue circles, left). The effect of photoactivation and region selected for kymograph generation (dashed white rectangle, right). Scale bar, 10 μm. **D.** Corresponding kymograph profiles of the photoactivated regions in bipolar and monopolar spindles used for quantification of the flux rates (red dotted lines highlight MT-flux slopes). Scale bars, 30 sec. **E, F**. Quantification of MT-flux in bipolar (E) and monopolar (F) spindles subjected to indicated treatments. Graphs represent MT flux of individual cells with mean ± SD. N (number of cells, number of independent experiments): - bipolar spindles: siControl (49, 5), STLC (31, 3), siKIF15 (39, 3), siKID (32, 3), siKIF4A (28, 3), siKIF2A (36, 3), siCLASPs (21, 2), GSK923295 (44, 3); - monopolar spindles: siControl (35, 3), siKIF15 (44, 3), siKID (38, 3), siKIF4A (44, 3), siKIF2A (38, 3), siCLASPs (12, 2), GSK923295 (33, 3). P-values were calculated using one-way ANOVA. n.s. - not significant, **** P ≤ 0.0001.

### KIF4A promotes MT-flux via its chromosome arm-based motor activity

Because of its strong contribution, we sought to characterize the molecular mechanisms by which the chromokinesin KIF4A mediates MT-flux. First, we examined the effect of KIF4A overexpression on MT-flux rates. To facilitate the measurement of dose response effects, we cloned KIF4A into an adenoviral vector to infect target cells with increasing virus titers (multiplicity of infection ratios (MOI)) (**Fig EV1B**). KIF4A overexpression resulted in increased MT-flux rates (**Fig EV1C**), supporting the hypothesis that KIF4A can act as a MT-flux driving force. Although RNAi and overexpression experiments clearly showed the importance of KIF4A for MT-flux, it remained to be tested how KIF4A contributed to this process. To address this question, we conditionally reconstituted KIF4A-depleted cells with close to physiological amounts of wild-type or mutant KIF4A (Sigl, Ploner et al., 2014) (**Fig 2A and B, and Movie EV3**). Using this approach, we designed an RNAi rescue experiment to assess whether RNAi resistant versions of wild-type KIF4A, the ATPase-dead K94A motor-mutant as well as a chromatin non-binding KIF4A mutant (ΔZip1) that was still able to bind to central spindle interpolar MTs (Wu & Chen, 2008) (**Fig 2A**), would restore MT-flux in KIF4A RNAi cells. While wild-type KIF4A successfully rescued the reduced MT-flux rates in KIF4A RNAi cells (0.53 +/− 0.15 μm/min, compared to 0.57 +/− 0.1 μm/min in uninduced cells and to 0.29 +/− 0.13 μm/min in KIF4A shRNA), neither the motor mutant (K94A) (0.33 +/− 0.14 μm/min, compared to 0.63 +/− 0.15 μm/min in uninduced cells), nor the chromatin-binding mutant (ΔZip1) (0.3 +/− 0.1 μm/min, compared to 0.63 +/− 0.13 μm/min in uninduced cells) were able to recover MT-flux rates (**Fig 2C**). These results indicate that both the motor activity and its localization on chromosome arms are essential for MT-flux (**Fig 2D**).

**Figure 2.**
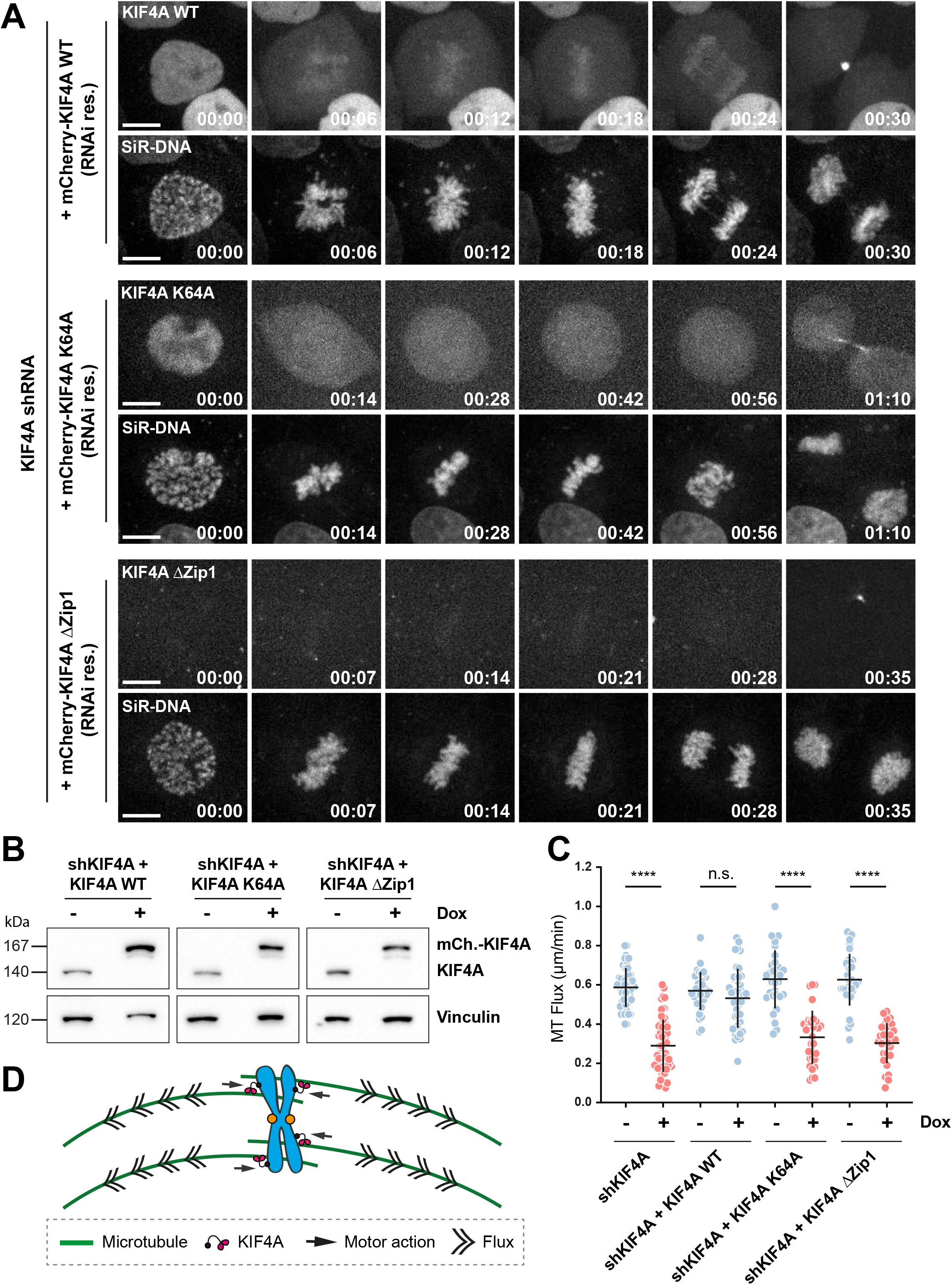
KIF4A promotes MT-flux via its chromosome arm-based motor activity. **A**. Representative spinning disk confocal live-cell imaging time series images of U2OS PA-GFP-α- tubulin cells conditionally co-expressing KIF4A shRNA and RNAi-resistant mCherry-Kif4A variants induced using doxycycline. Chromosomes were stained using SiR-DNA. Scale bars, 10 μm. Time, hour:min. **B.** Representative immunoblot of cell lysates obtained before and after doxycycline induction validating the efficiency of KIF4A shRNA construct and expression of the RNAi-resistant mCherry-KIF4A variants stained using anti-KIF4A antibody, with anti-vinculin used as a loading control. **C.** Quantification of MT-flux upon shRNA-mediated depletion of KIF4A alone or in combination with conditional expression of the RNAi-resistant mCherry-KIF4A variants. The bars in graph represent mean ± SD. N (number of cells, number of independent experiments): uninduced shKIF4A (48, 4), shKIF4A (60, 4), uninduced KIF4A WT (36, 5), shKIF4A + KIF4A WT (43, 5), uninduced KIF4A K64A (30, 4), shKIF4A + KIF4A K64A (32, 4), uninduced KIF4A ΔZip1 (32, 3), shKIF4A + ΔZip1 (30, 4). P-values were calculated using Student’s *t*-test. n.s. - not significant, **** P ≤ 0.0001. **D.** Model illustrating chromosome arms-localized KIF4A driving MT-flux.

Based on these data, our model supports that KIF4A at the chromosome arms interacts with non-KT-MTs and, by walking towards their plus-ends, it helps chromosomes to align at the spindle equator in metaphase. Once all the chromosomes have become aligned, KIF4A cannot push them any further due to equivalent forces applied from opposite spindle sides. However, KIF4A still remains processive on the interacting MTs, thereby providing the reactive force that promote the poleward flux (**Fig EV1D**).

To further elucidate the involvement of chromatin in MT-flux generation, we successfully established U2OS cells undergoing mitosis with unreplicated genomes (MUGs) (Brinkley, Zinkowski et al., 1988, O’Connell, Loncarek et al., 2008) (Brinkley et al., 1988, O’Connell et al., 2008) (**Fig 3A and C, EV3, and Movie EV4**), providing a way to test its contribution to MT-flux. In agreement with the data obtained using KIF4A mutants, MT-flux in bipolar MUG spindles was strongly reduced (0.31 +/− 0.13 μm/min, compared to 0.5 +/− 0.13 μm/min in control cells treated with caffeine alone) (**Fig 3B**), further highlighting the importance of chromatin as the locus of KIF4A activity relevant for MT-flux by interacting with non-KT-MTs.

**Figure 3.**
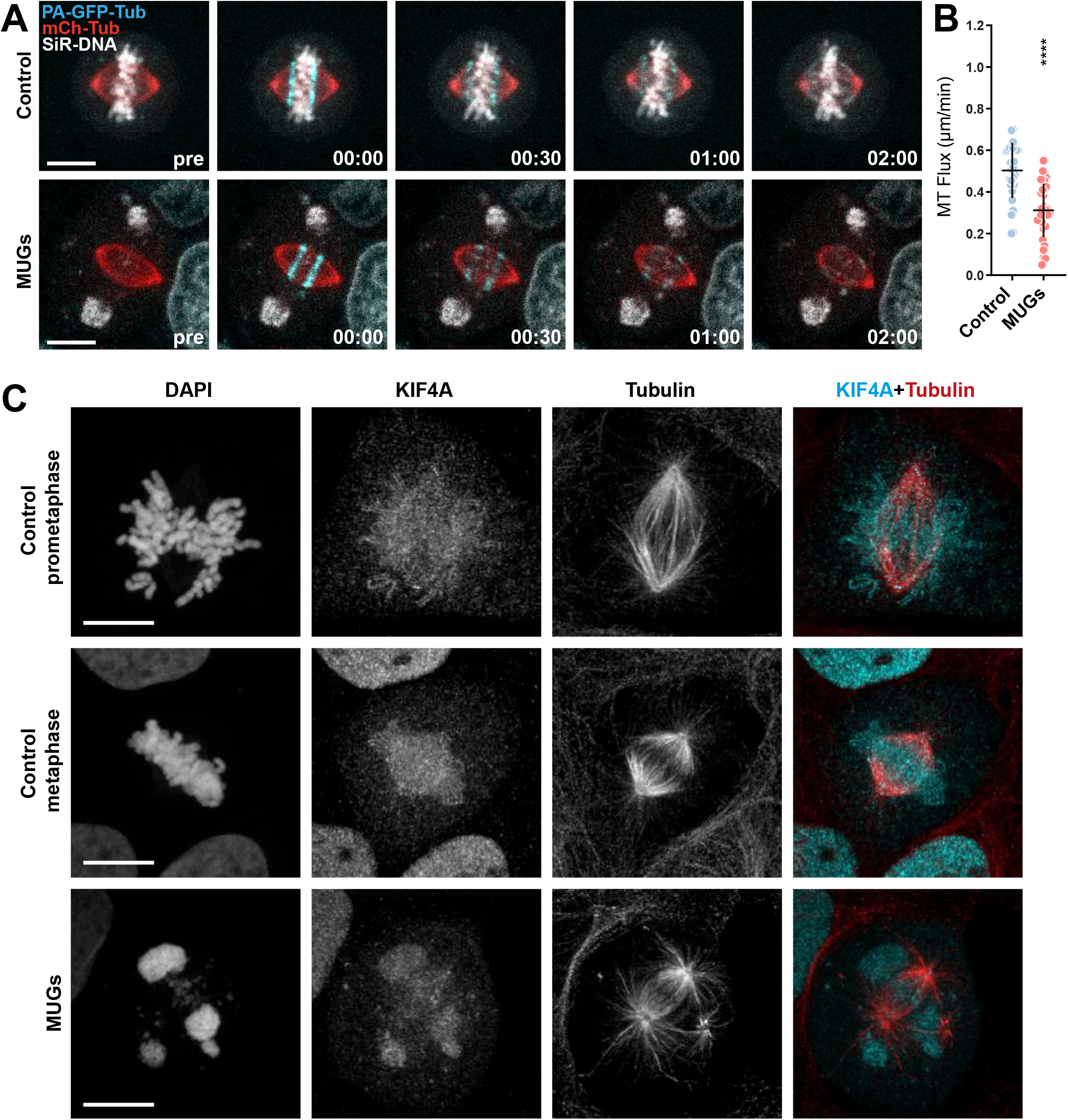
Bipolar spindles with expelled chromatin in MUGs exhibit reduced MT-flux. **A.** Representative spinning disk confocal time series of MT-flux in bipolar spindles in control cells and cells undergoing mitosis with unreplicated genomes (MUGs). U2OS cells stably co-expressing PA-GFP-α-tubulin (cyan) and mCherry-α-tubulin (red), labeled for chromosomes with SiR-DNA (grey), are shown, with 5 mM caffeine used as a control. Scale bar, 10 μm. Time, min:sec. **B.** Quantification of the MT-flux rates from indicated conditions. MT-flux values with mean ± SD are plotted. N (number of cells, number of independent experiments): Caffeine-only control (30, 3) and MUGs (34, 2). P-values from Student’s t test. **** P ≤ 0.0001. **C**. Representative point-scanning confocal maximum-intensity projected images of mitotic spindles in U2OS cells subjected to indicated conditions, immunostained with antibodies against KIF4A and α-tubulin. DNA was counterstained with DAPI. KIF4A in cyan and α-tubulin in red in merged image. Scale bar, 10 μm.

### MT-crosslinking proteins HSET and NuMA facilitate the distribution of MT-flux-dependent forces across the mitotic spindle

If our model based on KIF4A driving MT-flux from chromosome arms was correct, one would predict that the action of MT cross-linking molecules is required to transmit the fluxing forces from non-KT-MTs to KT-MTs, as proposed by the “coupled spindle” model (Matos et al., 2009). In order to test this hypothesis, we investigated the contribution of three well-established MT-crosslinking proteins to MT-flux, namely NuMA, kinesin-14/HSET and PRC1. NuMA is recruited to the spindle poles by the MT minus-end-directed motor Dynein to cross-link and focus MT minus-ends (Merdes, Ramyar et al., 1996, Radulescu & Cleveland, 2010). HSET is a MT minus-end-directed kinesin motor with the ability to crosslink and slide anti-parallel MTs, thereby generating an inward force within the spindle, similar to that proposed for Dynein (Mountain, Simerly et al., 1999). PRC1 is involved in the assembly of the central spindle during late mitosis (Mollinari, Kleman et al., 2002) and was proposed to bridge KT-MTs with interpolar MTs during spindle assembly (Kajtez, Solomatina et al., 2016, Polak, Risteski et al., 2017, Vukusic, Buda et al., 2017). We found that, while depletion of NuMA and HSET, individually or together, significantly reduced KT-MT-flux rates (0.48 +/− 0.13 μm/min, 0.36 +/− 0.13 μm/min, and 0.35 +/− 0.16, respectively) (**Fig 4A and B, EV2A, and Movie EV5**), depletion of PRC1 had no such effect (0.61 +/− 0.22 μm/min, compared to 0.61 +/− 0.22 μm/min in control cells) (**Fig 4B and EV2A**). Thus, KIF4A’s effect on MT-flux is independent of its PRC1-dependent localization at interpolar MTs (Zhu & Jiang, 2005).

**Figure 4.**
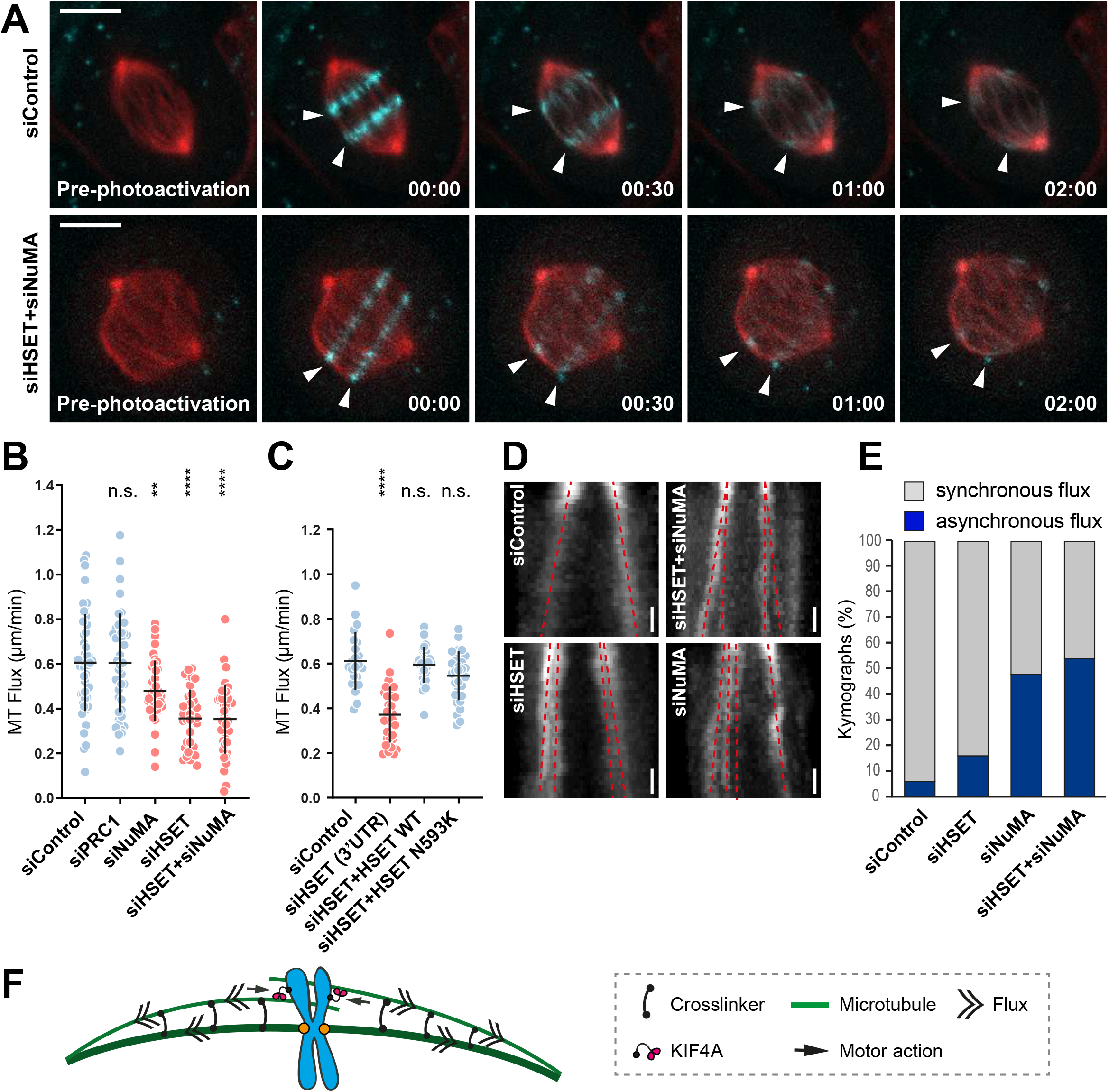
MT-crosslinking activities of NuMA and kinesin-14/HSET ensure the uniform distribution of MT-flux-dependent forces across the mitotic spindle. **A**. Representative spinning disk confocal live-cell image series of MT-flux in U2OS cells stably co-expressing PA-GFP-α- tubulin (cyan) and mCherry-α-tubulin (red) treated with indicated siRNAs. White arrowheads highlight poleward motion of the photoactivated regions due to MT-flux. Scale bars, 10 μm. Time, min:sec. **B.** Quantification of the impact of the MT-crosslinking proteins on MT-flux in U2OS PA-GFP/mCherry-α-tubulin cells transfected with respective siRNAs. Graph represent MT-flux values with mean ± SD. N (number of cells, number of independent experiments): siControl (49, 5), siPRC1 (45, 3), siNuMA (40, 3), siHSET (37, 3), siHSET + siNuMA (39, 3). P-values were calculated using one-way ANOVA. n.s., not significant, ** P ≤ 0.01, **** P ≤ 0.0001. **C**. Quantification of the impact of the motor activity of HSET on MT-flux in U2OS cells stably expressing mEOS-α-tubulin treated with control or HSET 3′UTR siRNAs in presence or absence of the respective RNAi-resistant GFP-HSET constructs. Graph represent MT-flux values with mean ± SD. N (number of cells, number of independent experiments): siControl (23, 3), siHSET 3′UTR (33, 3), siHSET 3′UTR + HSET WT (29, 5), siHSET 3′UTR + HSET N593K (30, 4). P-values were calculated using one-way ANOVA. n.s., not significant, **** P ≤ 0.0001. **D.** Representative kymographs of the photoactivated spindles from (B) displaying asynchronous flux motion (split of the two red dashed lines towards individual poles) upon RNAi-mediated depletion of MT-crosslinkers. **E.** Percentage of cells with asynchronous flux movements calculated from (B). **F.** Model illustrating the role of MT-crosslinking activities of NuMA and HSET in uniform distribution of poleward forces across the mitotic spindle.

In order to dissect whether the contribution of HSET for MT-flux depended either on its MT crosslinking ability, or on its motor activity, we designed a rescue experiment in which we individually expressed WT HSET and a non-processive motor-mutant that is still able to crosslink MTs (N593K) (Cai, Weaver et al., 2009). Because our HSET constructs were GFP-tagged and could thus interfere with the PA-GFP tubulin signal, we used a U2OS cell line stably expressing green-red photoconvertible mEOS-tubulin (Wandke et al., 2012) (**Fig EV2B**). By monitoring flux rates after depletion of endogenous HSET by 3’UTR-targeting siRNAs (0.37 +/− 0.13 μm/min, compared to 0.61 +/− 0.13 μm/min in control cells), we observed that both WT HSET and the motor-mutant successfully ensured normal MT-flux (0.59 +/− 0.1 μm/min and 0.55 +/− 0.11 μm/min, respectively) (**Fig 4C and EV2C**). This implies that HSET contributes to KT-MT-flux by coupling KT-MTs to flux-driving non-KT-MTs (**Fig 4F**). Interestingly, the kymograph profiles of photoactivated spindles depleted of NuMA and HSET revealed frequent occurrence of asynchronous flux movements (6% in control cells, 16% in HSET RNAi, 48% in NuMA RNAi and 54% in HSET+NuMA RNAi) (**Fig 4D and E**), which may be explained by a more frequent disengagement between flux-driving MTs and KT-MTs, indicating weakened mechanical coupling. Altogether, these data suggest that MT-crosslinking activities of NuMA and kinesin-14/HSET are required to transmit the MT-flux forces generated by KIF4A on non-KT-MTs to KT-MTs, thereby promoting a uniform distribution of poleward forces across the mitotic spindle (**Fig 4F**).

### CENP-E is the predominant driver of MT-flux in early prometaphase

Another prediction of the “coupled spindle” model is that KT-MTs flux at slightly slower rates compared to non-KT-MTs due to imperfect coupling (LaFountain, Cohan et al., 2004, Maddox et al., 2003, Matos et al., 2009). To test this prediction in human cells, we measured flux rates after perturbation of end-on KT-MT attachments by depletion of NDC80/HEC1, the main MT-anchoring protein at KTs (Cheeseman, Chappie et al., 2006, DeLuca, Moree et al., 2002, McCleland, Gardner et al., 2003, Wei, Al-Bassam et al., 2007, Wigge & Kilmartin, 2001) (**Fig EV3A)**. Indeed, MTs in these cells fluxed faster (0.91 +/− 0.2 μm/min) (**Fig 5A and B, and Movie EV6**), suggesting that stable end-on KT-MT attachments work as flux brakes, in agreement with the “coupled spindle” model. Remarkably, in contrast to its effect in cells with established end-on attachments (**Fig 1A and E**), depletion of KIF4A did not compromise the flux rate after NDC80 RNAi (**Fig 5B and EV3A**). Thus, KIF4A is dispensable for MT-flux prior to the establishment of end-on KT-MT attachments (**Fig 4F and EV1D**).

**Figure 5.**
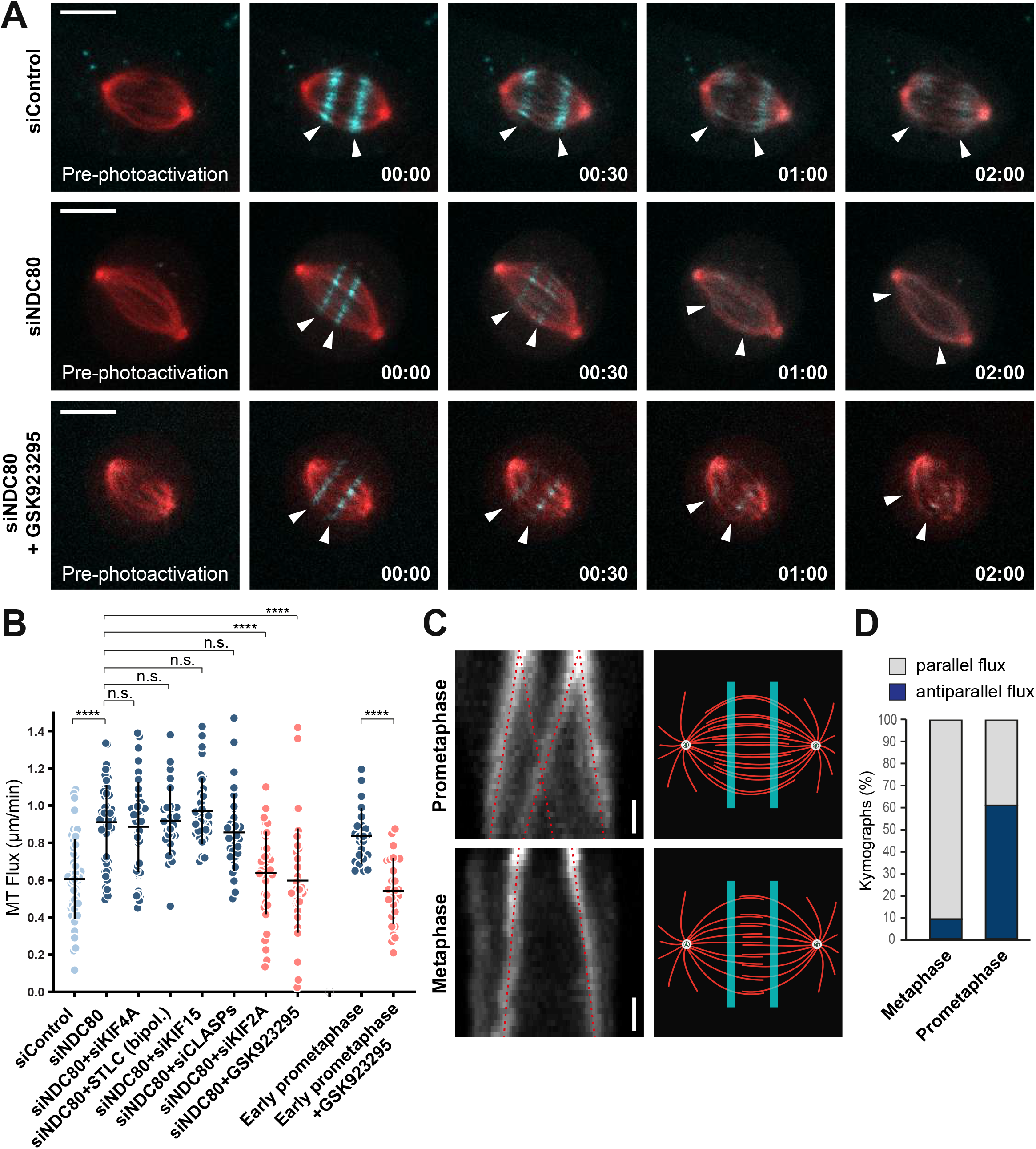
CENP-E is the predominant driver of MT-flux in early prometaphase. **A**. Representative spinning disk confocal live-cell time series of MT-flux in U2OS cells stably co-expressing PA-GFP-α-tubulin (cyan) and mCherry-α-tubulin (red) treated with indicated conditions. White arrowheads follow the poleward motion of the photoactivated regions due to MT-flux. Scale bars, 10 μm. Time, min:sec. **B.** Quantification of the impact of stable end-on KT-MT attachments on MT-flux in NDC80-depleted cells under indicated conditions and in early prometaphase cells with and without the CENP-E inhibitor GSK923295. MT-flux values are plotted with mean ± SD. N (number of cells, number of independent experiments): siControl (49, 5), siNDC80 (59, 6), siNDC80 + siKIF4A (41, 3), siNDC80 + STLC (bipol.) (28, 3), siNDC80 + siKIF15 (36, 4), siNDC80 + siCLASPs (33, 3), siNDC80 + siKIF2A (38, 4), siNDC80 + GSK923295 (38, 3), early prometaphase (23, 3), early prometaphase + GSK923295 (28, 3). P-values were calculated using Student’s *t*-test or one-way ANOVA. n.s., not significant, **** P ≤ 0.0001. **C.** Kymograph profiles (left) highlighting that the photoactivated regions in prometaphase (upper panel) split and move towards both poles, whereas metaphase spindles (lower panel) flux uniformly towards individual poles (indicated by red dotted lines). Illustration of the photoactivated regions (right) in prometaphase and metaphase spindles, highlighting that interpolar MTs overlap over a broader region in early mitosis, becoming focused at the spindle equator in metaphase. Scale bar, 30 sec **D.** Percentage of cells with parallel and antiparallel flux movements during metaphase and prometaphase. N (number of cells, number of independent experiments): prometaphase (23, 3), metaphase (22, 3).

To investigate what drives MT-flux before end-on KT-MT attachments are established, we further examined the potential contributions of EG5 and KIF15 (**Fig 5B and EV3A**). Similar to the effect of KIF4A depletion, neither EG5 inhibition nor depletion of KIF15 reduced the flux rate after NDC80 RNAi. Since CENP-E inhibition showed a negative impact on flux rate only in monopolar spindles that contain less end-on KT-MT attachments (Shrestha & Draviam, 2013) (**Fig 1E and F**), we inhibited CENP-E in NDC80-depleted cells. Unlike its knockdown, chemical inhibition of CENP-E keeps the inactive motor at KTs, allowing its interaction with CLASPs, thus excluding any potential indirect effects (Logarinho et al., 2012, Maffini et al., 2009). Strikingly, CENP-E inhibition in NDC80-depleted cells strongly decreased the flux rate (0.6 +/− 0.28 μm/min, compared to 0.91 +/− 0.2 μm/min in NDC80 RNAi) (**Fig 5A and B, and Movie EV6)**, similar to the effect of NDC80 co-depletion with KIF2A (0.64 +/− 0.22 μm/min) (**Fig 5B and EV3A**). However, co-depletion of NDC80 and CLASPs did not reduce the flux rate (0.86 +/− 0.21 μm/min) compared to NDC80 knockdown (**Fig 5B and EV3A**), demonstrating that CLASPs contribute to MT-flux exclusively by maintaining the growth of KT-MTs (Girao, Okada et al., 2020).

Taken together, these data suggest that CENP-E promotes MT-flux in early prometaphase, prior to the establishment of stable end-on KT-MT attachments. To directly test this, we first compared flux rates between early and late prometaphase/metaphase and found that MT-flux in early prometaphase was indeed faster (0.84 +/− 0.15 μm/min and 0.61 +/− 0.22 μm/min, respectively) (**Fig 5B and EV4A**), consistent with the increased flux found in NDC80-depleted cells. Subsequent measurement of MT-flux in CENP-E-inhibited early prometaphase cells showed a significant decrease in the flux rate (0.54 +/− 0.18 μm/min) (**Fig 5B**), comparable to the one observed upon CENP-E inhibition in NDC80-depleted cells. We conclude that CENP-E motor activity is required for MT-flux prior to the establishment of end-on KT-MT attachments, independently from its role in CLASPs recruitment to KTs. Noteworthy, unlike in metaphase, photoactivated marks in early prometaphase showed splitting and moved simultaneously towards both poles (61% in prometaphase, compared to 9% in metaphase) (**Fig 5C and D, EV4A, and Movie EV7**), as it was observed in in-vitro-assembled *Xenopus* egg extract spindles (Sawin & Mitchison, 1991). This strongly suggests that interpolar MTs overlap over a broader region in early mitosis, but then become focused at the spindle equator as cells reach metaphase. The broad distribution of antiparallel MTs in early mitosis coincides with CENP-E localization at laterally attached KTs during “prometaphase rosette” configuration (Itoh, Ikeda et al., 2018) (**Fig 6A and B, and EV3B**), and is consistent with the localization of the antiparallel MT-crosslinker PRC1, which also covers a wider region in prometaphase (**Fig EV4B**).

**Figure 6.**
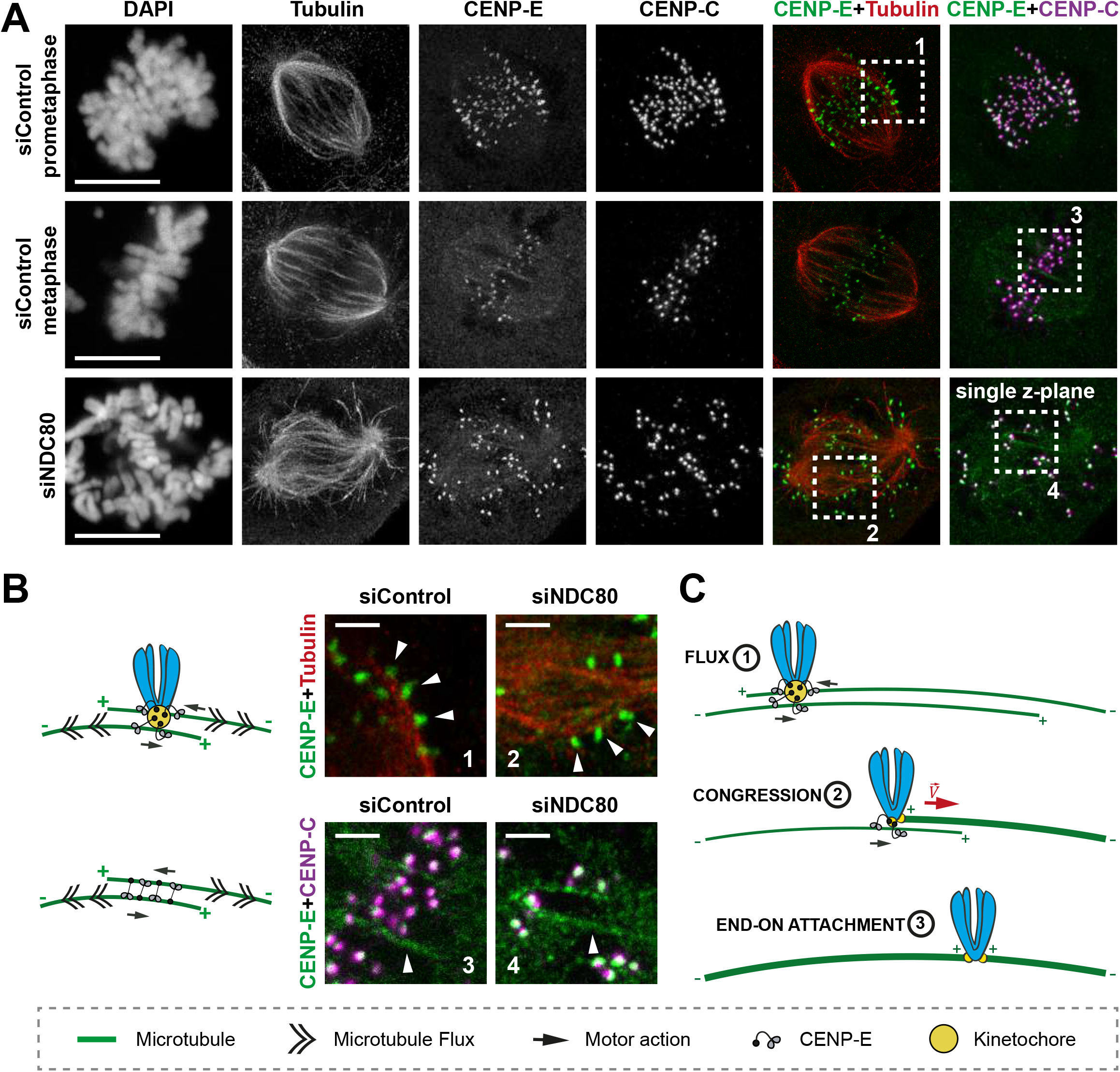
Analysis of CENP-E localization using super-resolution coherent-hybrid STED (CH-STED) microscopy. **A.** Representative maximum-intensity projected coherent-hybrid STED (CH-STED) images of mitotic spindles in U2OS cells treated with control and NDC80 siRNAs and stained for DNA (DAPI), KTs (CENP-C, magenta), α-tubulin (red) and CENP-E (green). Scale bar, 10 μm. **B.** Model (left) illustrating potential mechanisms for CENP-E mediated MT-flux forces either through its KT- or antiparallel MT-based localization and sliding. Zoomed insets (right) from (a) highlight CENP-E localized at KTs laterally interacting with MTs (white arrowheads in **1**, **2**), and CENP-E localized at interpolar MTs (white arrowheads in **3**, **4**), in control and NDC80-depleted mitotic cells. Scale bars, 2 μm. **C.** Model illustrating MT-flux driven by CENP-E and its potential impact during early stages of mitosis. CENP-E at laterally attached KTs interacts with interpolar MTs and slides them apart until MT-plus-ends reach the KT and become end-on attached, supporting chromosome congression and transition from lateral to end-on KT-MT attachments. See text for more details.

To gain a deeper mechanistic insight into how CENP-E promotes MT-flux in early mitosis, we used super-resolution coherent-hybrid STED (CH-STED) microscopy (Pereira, Sousa et al., 2019) to determine CENP-E localization (**Fig 6A and B**). Surprisingly, apart from its well-established localization at KTs (**Fig 6A and B, and EV3B**) (Kapoor et al., 2006, Schaar et al., 1997, Wood et al., 1997, Yen et al., 1991) and antiparallel midzone MTs during late anaphase and telophase (Yen et al., 1991), we found that CENP-E also localized at interpolar MTs, both in control and NDC80-depleted mitotic cells (**Fig 6A and B**). These observations raised two possible explanations for how CENP-E could drive MT-flux prior to the establishment of end-on KT-MT attachments. Given that CENP-E localizes at the expanded fibrous corona of unattached outer KTs during early mitosis (**Fig 6A and B, and EV3B**), KT-localized CENP-E might interact with antiparallel interpolar MTs, sliding them apart towards the poles (**Fig 6B**). Another possibility would be that CENP-E directly crosslinks antiparallel MTs and slides them apart (**Fig 6B**). Since CENP-E contributes to MT-flux in STLC-treated monopolar spindles (**Fig 1F**), which lack antiparallel MTs, but contain a significant fraction of laterally attached KTs (Shrestha & Draviam, 2013) with CENP-E (**Fig EV3B**), these data favor the first scenario, without excluding a contribution from direct MT crosslinking of antiparallel MTs in bipolar spindles.

### Combined action of EG5 and KIF15 support MT-flux driving activities of CENP-E and KIF4A

Although we identified CENP-E and KIF4A as drivers of MT-flux in prometaphase and metaphase, respectively, functional inactivation of either of these two motors resulted in only around 50% decrease in MT-flux rate, indicating the presence of other important contributor(s). Therefore, we tested whether EG5 and KIF15 play a synergistic role in supporting MT-flux. Importantly, in contrast to their individual functional inactivation, which did not affect MT-flux rates (**Fig 1E**), inhibition of EG5 in KIF15-depleted cells resulted in a significant decrease of MT-flux in bipolar spindles (0.43 +/− 0.11 μm/min, compared to 0.61 +/− 0.22 μm/min in controls), as well as in NDC80-depleted spindles (0.51 +/− 0.13 μm/min, compared to 0.91 +/− 0.2 μm/min in NDC80 RNAi only) (**Fig 7B and C**), suggesting that these two tetrameric antiparallel MT crosslinking motors play redundant roles, similar to their role in bipolar spindle assembly (Tanenbaum et al., 2009). Therefore, we examined whether MT-flux is driven by cooperative actions between CENP-E, EG5 and KIF15 prior to the establishment of stable end-on KT-MT attachments. Strikingly, triple inactivation of CENP-E, EG5 and KIF15 resulted in virtually no flux both in NDC80-depleted- (0.1 +/− 0.09 μm/min, compared to 0.91 +/− 0.2 μm/min in NDC80 RNAi only) and in early prometaphase cells (0.07 +/− 0.1 μm/min, compared to 0.84 +/− 0.15 μm/min in control early prometaphase cells) (**Fig 7A and Movie EV8**). Next, we tested whether EG5 and KIF15 also cooperate with KIF4A to drive MT-flux upon the establishment of stable end-on KT-MT attachments. Triple inactivation of KIF4A, EG5 and KIF15 in late prometaphase/metaphase cells led to a much stronger reduction in MT-flux, compared to either KIF4A-depletion or co-inactivation of EG5 and KIF15 (0.15 +/− 0.1 μm/min, compared to 0.43 +/− 0.1 μm/min in STLC+KIF15 RNAi and 0.32 +/− 0.14 in KIF4A RNAi) (**Fig 7C and Movie EV8**). Altogether, these data demonstrate that mitotic MT-flux is driven by the coordinated action of 4 kinesins.

**Figure 7.**
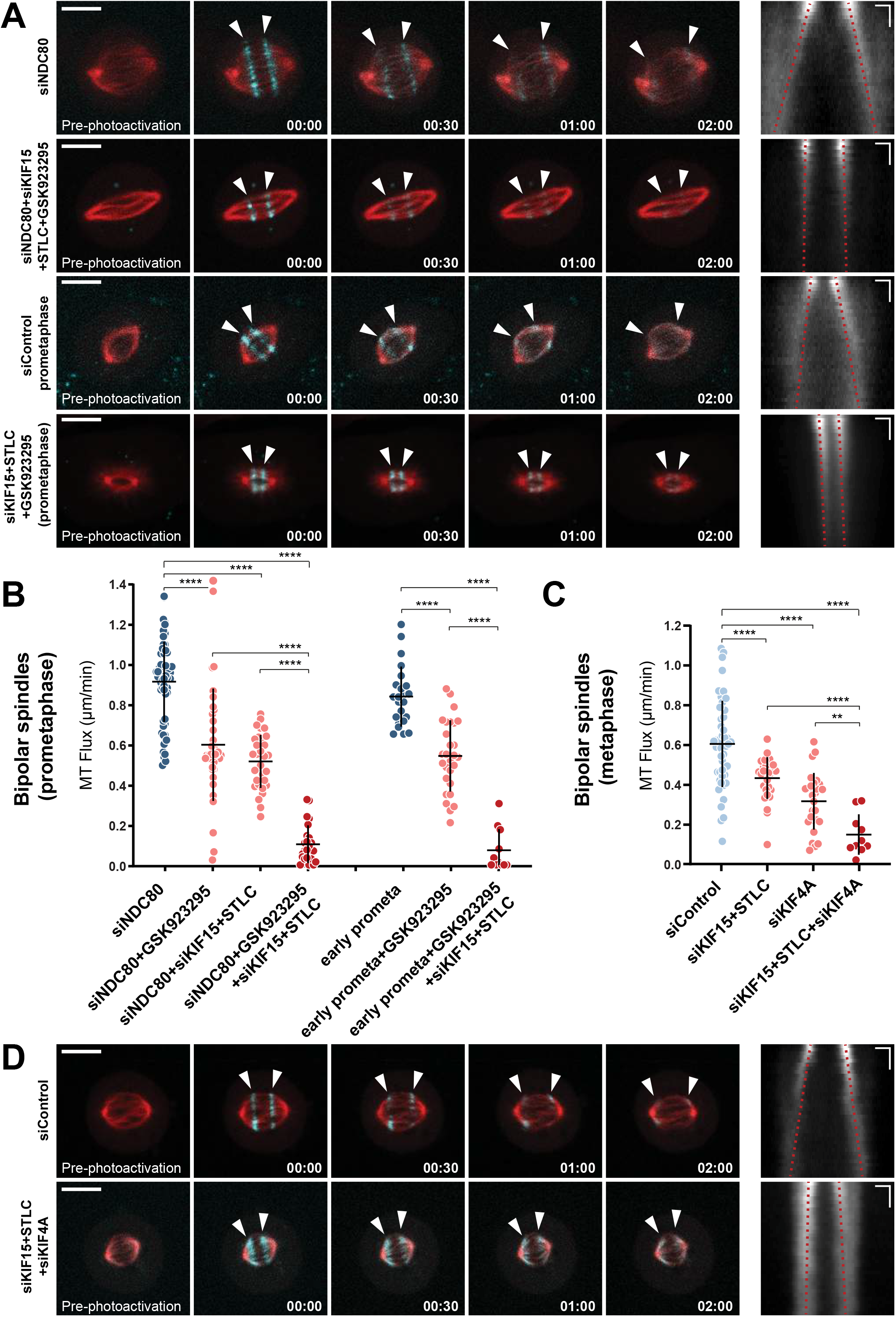
Combined action of EG5 and KIF15 support MT-flux driving activities of CENP-E and KIF4A. **A.** Representative spinning disk confocal time series of U2OS cells stably co-expressing PA-GFP-α-tubulin (cyan) and mCherry-α-tubulin (red), with indicated treatments (left). White arrowheads highlight poleward flux of the photoactivated regions. Scale bar, 10 μm. Time, min:sec. Right panels display corresponding kymographs of the photoactivated spindles used for quantification of the flux rates (red dotted lines highlight MT-flux slopes). Scale bars, 2 μm (horizontal) and 30 sec (vertical). **B, C**. Quantification of MT-flux in prometaphase (B) and metaphase (C) bipolar spindles subjected to indicated treatments. Graphs represent MT flux of individual cells with mean ± SD. siNDC80, siNDC80 + GSK923295, early prometaphase, early prometaphase + GSK923295, siControl, and siKIF4A data are from Figure 1E and 5B. N (number of cells, number of independent experiments): siNDC80 + KIF15 + STLC (33, 5), siNDC80 + GSK923295 + KIF15 + STLC (25, 3), early prometaphase + GSK923295 + KIF15 + STLC (11, 2), siKIF15 + STLC (29, 3), siKIF15 + STLC + siKIF4A (10, 3). P-values were calculated using one-way ANOVA. ** P ≤ 0.01, **** P ≤ 0.0001. **D.** Representative spinning disk confocal time series of U2OS cells stably co-expressing PA-GFP-α-tubulin (cyan) and mCherry-α-tubulin (red), with indicated treatments (left) and corresponding kymographs (right) as described in (A).

### MT poleward flux regulates spindle length in response to MCAK-mediated depolymerization of KT-MTs

Similar to what was observed upon CLASPs depletion (Maffini et al., 2009), we observed substantial reduction of spindle length under certain conditions that resulted in highly attenuated MT-flux rates (**Fig 7A and D**). To investigate the relationship between MT-flux and spindle length, we analyzed the correlation between altered MT-flux rates and spindle sizes from different treatments. We found that MT-flux rate and spindle size showed a strong positive correlation overall (Pearson correlation coefficient, r = 0.73; p value = <0.0001) (**Fig 8A and Table EV1**). Importantly, we noticed that spindle size did not dramatically change in NDC80-depleted cells with reduced flux rates, where no correlation between MT-flux rate and spindle size was observed (Pearson correlation coefficient, r = 0.68; p value = 0.2063) (**Fig 8C**). Accordingly, when we analyzed the correlation between MT-flux and spindle size in cells containing mainly stable end-on KT-MT attachments that displayed reduced flux, the correlation between MT-flux rate and spindle size was strong (Pearson correlation coefficient, r = 0.87; p value = 0.0099) (**Fig 8B**).

**Figure 8.**
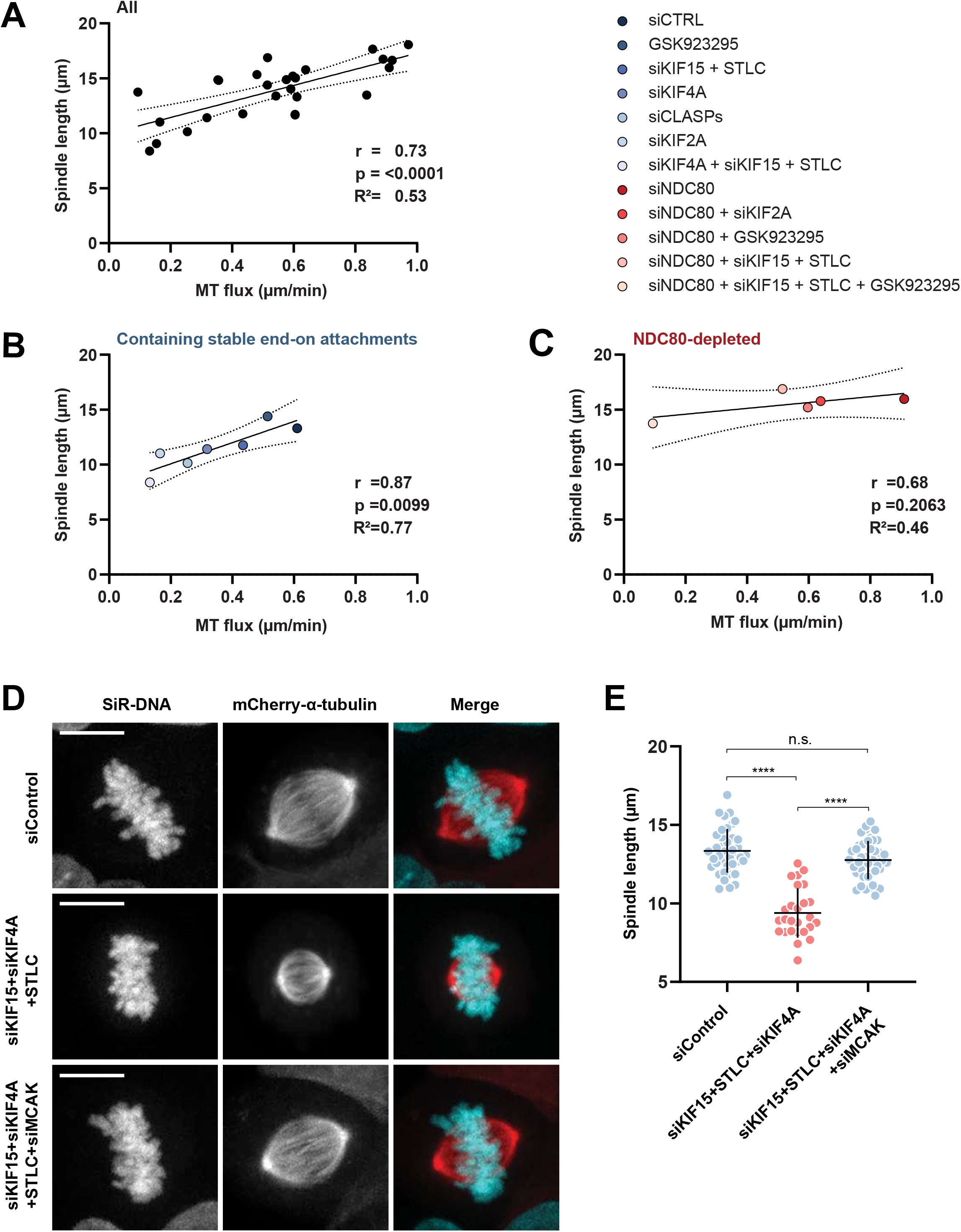
MT poleward flux regulates spindle length in response to MCAK-mediated depolymerization of KT-MTs. **A-C.** Graph depicting a direct correlation between MT-flux and mitotic spindle where mean spindle length is plotted over mean MT-flux rates for all the indicated conditions (A), mitotic conditions containing stable end-on attachments (B), and NDC80 depleted conditions which lack stable end-on attachments (C). Solid and dotted lines represent the linear regression and 95% confidence interval respectively. Pearson correlation coefficient (r), coefficient of determination (r^2^) and corresponding p values are indicated. **D.** Representative images from spinning disk confocal live-cell time series of U2OS PA-GFP-α-tubulin/mCherry-α-tubulin cells where DNA is labelled with SiR-DNA with the indicated treatments. Microtubules are shown in red and DNA in cyan in the merged image. Scale bar: 10 μm. **E.** Quantification of spindle lengths at metaphase following indicated treatments. Graph represents the spindle length of individual cells with mean ± SD. N (number of cells, number of independent experiments): siControl (41, 3), siKIF15 + STLC + siKIF4A (26, 3), siKIF15 + STLC + siKIF4A + siMCAK (43, 3). P-values were calculated using one-way ANOVA. n.s. - not significant, **** P ≤ 0.0001.

These data suggest that MT-flux regulates spindle size in response to the formation of stable end-on KT-MT attachments (**Fig EV5A**). Previously it was shown that although depletion of MCAK (kinesin-13-family member) alone did not affect MT-flux rate, its co-depletion with KIF2A not only strongly reduced MT-flux rate but also restored spindle bipolarity and size in KIF2A depleted cells [11]. Because simultaneous inactivation of KIF4A, EG5 and KIF15 in late prometaphase/metaphase cells led to much shorter spindles (**Fig 7D, 8A and B, Table EV1**), we tested whether MT-flux regulates spindle size by preventing MCAK-mediated depolymerization of k-fibres. Strikingly, we found that spindle shortening after triple inactivation of KIF4A, EG5 and KIF15 was fully rescued upon co-depletion of MCAK (12.76 +/− 1.22 μm compared to 9.39 +/− 1.57 μm in triple-inactivated cells and 13.35 +/− 1.39 μm in controls) (**Fig 8D and E, and EV5B**). In addition to its effect on spindle size, MCAK co-depletion significantly reduced spindle collapsing in bipolar triple-inactivated cells (**Fig EV5C**). Overall, these data indicate that MT-flux regulates spindle size by counteracting MCAK-driven MT-depolymerizing activity at KTs.

## Discussion

Mitotic spindle MTs remain highly dynamic even when they maintain a certain length, which is associated with their poleward fluxing and balancing their polymerization and depolymerization at their plus- and minus-ends, respectively. However, the functional role and regulation of MT poleward flux has remained elusive.

Our study not only establishes the long-sought molecular mechanisms underlying MT-flux, but also explains its role in maintaining spindle size upon the establishment of stable end-on KT-MT attachments and thus provides evidence that MT-flux is not only an epiphenomenon of MT-dynamics but an important requirement for spindle function. Altogether, our findings support a model in which MT-flux in human cells is driven by the coordinated action of four kinesins. Accordingly, interpolar MTs are slid apart by the collaborative activity of the tetrameric MT-crosslinking motors EG5 and KIF15, supported by sequential contribution of CENP-E at KTs in prometaphase and KIF4A on chromosome arms in metaphase (**Fig 9**).

**Figure 9.**
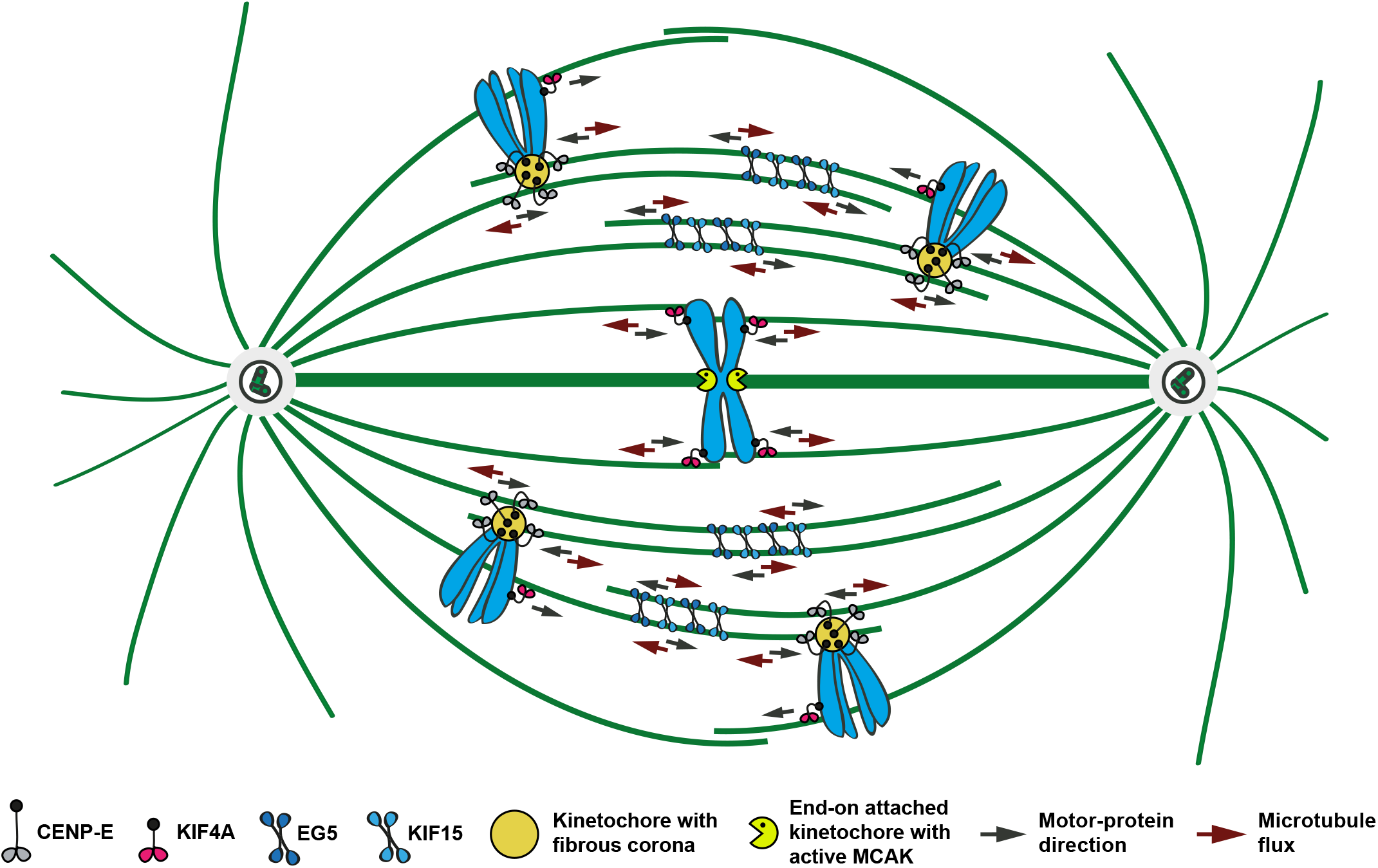
Model illustrating CENP-E- and KIF4A-dependent MT poleward flux driving forces. In our sequential MT-flux model, CENP-E at laterally attached KTs during early mitosis interacts with antiparallel MTs and slides them apart. Once chromosomes are aligned and CENP-E is partially stripped away from the KTs, KIF4A takes over promoting MTs to flux towards the poles.

In early mitosis, CENP-E at laterally attached KTs interacts with antiparallel interpolar MTs during congression and slides them apart until MT-plus-ends reach the KT and become end-on attached (**Fig 6C and Fig 9**). At this stage, CENP-E driven MT-sliding might be used to support chromosome congression when CENP-E uses laterally attached MTs as tracks towards the cell equator, supported by MT-depolymerization at the end-on attached sister KT (**Fig 6C and Fig 9**). During congression, end-on attachment is also established at the sister KT (**Fig 6C and Fig 9**), consistent with a role for CENP-E in the transition from lateral to end-on KT-MT attachments (**Fig 6C and Fig 9**) (Chakraborty, Tarasovetc et al., 2019, Shrestha & Draviam, 2013, Sikirzhytski, Renda et al., 2018). Upon bi-orientation, a large amount of CENP-E is stripped away from the KTs (Howell, McEwen et al., 2001) and our data indicates that KIF4A takes over at this stage to compensate for the weakened CENP-E contribution (**Fig 4F and EV1D**). Because KIF4A cannot move the chromosomes any further upon completion of chromosome congression due to equivalent opposite forces, KIF4A causes non-KT-MTs to flux towards the poles (**Fig 9 and EV1D**).

Interestingly, although CENP-E inhibition prior to establishment of end-on KT-MT attachments, as well as KIF4A depletion in metaphase, strongly affected MT-flux, the reduction was not complete (**Fig 1E and Fig 5B**). This was explained by the fact that simultaneous abrogation of the functions of EG5 and KIF15, together with CENP-E inhibition in early mitosis and KIF4A depletion in metaphase, respectively, resulted in virtually no flux (**Fig 7A-D**), demonstrating that MT-flux is driven by these four kinesins in human cells and further exposing the collaborative nature of MT-flux.

During early mitosis, CDK1 and PLK1 phosphorylate PRC1, keeping it mostly inactive (Hu, Ozlu et al., 2012, Zhu, Lau et al., 2006). At anaphase onset PRC1 is activated by dephosphorylation and brought to the midzone MTs by KIF4A (Zhu & Jiang, 2005). However, recent studies proposed PRC1 to bridge KT-MTs with interpolar MTs already during spindle assembly (Kajtez et al., 2016, Polak et al., 2017), opening the possibility that a pool of KIF4A co-localizes with PRC1 at interpolar MTs even before anaphase. Nevertheless, our KIF4A mutants- and PRC1 RNAi-based data revealed that the contribution of KIF4A to MT-flux depends on its motor activity and its localization on chromosome arms (**Fig 2A and C**), but not on its association with spindle midzone MTs (**Fig 2A and C, and 4B**).

Since KIF4A is not localized at KTs, the MT fluxing force generated at the chromosome arms has to be somehow transmitted to KT-MTs. We showed that MTs fluxed faster in NDC80-depleted cells that lack KT-MT end-on attachments, indicating that stable KT-MT end-on attachments normally work as flux brakes (**Fig 5A and B**), being in line with earlier observations in *Xenopus* egg extracts that, likely due to a drag, KT-MTs flux slightly slower than the neighboring interpolar MTs (Maddox et al., 2003), and that MT-flux is lost in spindles containing only KT-MTs (Mitchison, Maddox et al., 2004). This is also consistent with the “coupled spindle” model, in which fluxing non-KT-MTs transmit force to “passive” KT-MTs via MT-crosslinking (Matos et al., 2009). In agreement, we found that depletion of the MT-crosslinking proteins NuMA and HSET resulted in asynchronous MT-flux within different spindle MT subpopulations (**Fig 4D and E**). Thus, MT-crosslinking molecules ensure the uniform distribution of MT-flux-dependent forces across the mitotic spindle, enabling KT-MTs to flux due to coupling with non-KT-MTs (**Fig 4F**). Interestingly, chromosomes were shown to be dispensable for MT-flux in *Xenopus* egg extracts (Sawin & Mitchison, 1994), arguing against the role of chromatin in MT-flux driving. However, because the system lacked kinetochores as well (Sawin & Mitchison, 1994), it is reasonable that MT-flux depends on chromatin-based KIF4A only in the presence of stable end-on KT-MT attachments, as supported by the data presented in this study (**Fig 5B**).

Several studies implied that MT minus-end depolymerization is a response to flux, rather than its main driving force (Dumont & Mitchison, 2009, Maiato et al., 2004, Matos et al., 2009, Waterman-Storer et al., 1998). Our data here demonstrate that CLASPs are dispensable for MT-flux in the absence of end-on KT-MT attachments (**Fig 5B**). These findings suggest that MT-polymerizing activity of CLASPs and MT-depolymerizing activity of KIF2A, supported by MT-severing enzymes, whose depletion also led to a partial reduction in MT-flux rates (Jiang et al., 2017, Zhang, Rogers et al., 2007), act as MT-flux governing elements rather than its driving force. In addition, our study shows that although MT-flux rate is in general positively correlated with spindle size, this correlation is lost in cells without stable end-on KT-MT attachments. Because MCAK depletion fully restored spindle size in metaphase cells with reduced MT-flux, we conclude that the role of MT-flux in human cells is to regulate mitotic spindle size in response to KT-based MCAK activity. This explains why MT-flux becomes dispensable to regulate spindle length in the absence of either MCAK (Ganem et al., 2005) or stable end-on KT-MT attachments (**Fig 8A and C**), and why overexpression of MCAK leads to substantially shorter spindles (Liao, Rajendraprasad et al., 2019).

Altogether, our study demonstrates that MT mitotic flux in human cells is driven by the coordinated action of four kinesins, and is required to maintain mitotic spindle size by counteracting MCAK-mediated MT-depolymerizing activity at KTs.

## Materials and Methods

### Cell culture, cloning and plasmids

All cell lines were grown in Dulbecco’s Modified Eagle Medium (DMEM) supplemented with 10% fetal bovine serum (FBS; Thermo Fisher Scientific) at 37°C in humidified conditions with 5% CO2. U2OS photoactivatable GFP (PA-GFP)-α-tubulin (Ganem et al., 2005) cell line expressing stable inducible short hairpin RNA (shKIF4A) targeting KIF4A sequence 5’-GCAAGATCCTGAAAGAGAT-3’ was generated using a multipurpose GATEWAY-based lentiviral tetracycline-regulated conditional RNAi system (GLTR) using pENTR-THT-III (Addgene plasmid #55791) and pGLTR-X-Puro (Addgene plasmid #58246) plasmids (Pfeiffenberger, Sigl et al., 2016, Sigl et al., 2014). Stable inducible U2OS PA-GFP-α-tubulin KIF4A shRNA cell lines conditionally expressing RNAi-resistant mCherry-KIF4A variants were generated by lentiviral infection followed by clonal selection with antibiotics. The RNAi-resistant KIF4A and K94A motor mutant were generated using site-directed mutagenesis. RNAi targeting sequence 5’-GCAAGATCCTGAAAGAGAT-3’ was mutated into 5’-GCAAGATATTAAAAGAGAT-3’ (mutated nucleotides are underlined). Overlap extension PCR was used to generate RNAi-resistant KIF4A ΔZip1 mutant (Δ757-778) with KIF4A FWD: 5’-AAGGAAAAAAGCGGCCGCCATGAAGGAAGAGGTGAAGGGA-3’, KIF4A REV: 5’- CCGGATATCGTGGGCCTCTTCTTCGATAG-3’, KIF4AΔZip1 FWD: 5’- AACGAAATTGAGGTTATGGTCAGTGCTCAACTCAAAGAAAAAAAGGAA-3’, and KIF4AΔZip1 REV: 5’-TTCCTTTTTTTCTTTGAGTTGAGCACTGACCATAACCTCAATTTCGTA-3’. For Gateway cloning compatible N-terminal mCherry tagging of KIF4A, pENTR4-mCherry plasmid was generated by PCR amplification of the mCherry tag from pmCherry-C1 using the following primers mCh FWD: 5’- CGCGTCGACATGGTGAGCAAGGGCGAGGA-3’ and mCh REV: 5’-ACGCGGATCCCTTGTACAGCTCGTCCATGC-3’ and ligating it into pENTR4 no ccDB (gift from E. Campeau, Addgene plasmid #17424) plasmid (Campeau, Ruhl et al., 2009). The amplified PCR products of KIF4A variants were first cloned into pENTR4-mCherry, and then sub-cloned into pLenti-CMV/TO-DEST (gift from E. Campeau, Addgene plasmid #17291) (Campeau et al., 2009) by LR recombination (Invitrogen) according to manufacturer’s instructions. Production and expression of adenovirus encoding mCherry/mCherry-KIF4A was performed using pAd/CMV/V5-DEST Gateway Vector Kit (Thermo Fisher Scientific) according to manufacturer’s instructions. The plasmids and corresponding mutations were verified by sequencing.

For transient transfection experiments, all cDNA constructs were transfected (0.5 to 1μg) for 24 h using Lipofectamine 2000 (Thermo Fisher Scientific) according to the manufacturer’s protocol. For rescue experiments, GFP-HSET and GFP-HSET N593K plasmids (gift from C. Walczak, Indiana University School of Medicine, USA) (Cai et al., 2009) were used.

### Drug treatments

To induce monopolar spindles by inhibiting Eg5, 5 μM S-Trytil-L-Cystein (STLC; Santa Cruz) was added to the medium 20 min before live-cell imaging, or 5 h before fixation. CENP-E was inhibited by adding 200 nM GSK-923295 (Sigma-Aldrich) 20 min prior to imaging. DNA and microtubule cytoskeleton were labelled with 20 nM SiR-DNA and 20 nM SiR-tubulin (Spirochrome) respectively. To inhibit MT-flux, 10 nM taxol (LC laboratories) was added to the medium for 30 min before live-cell imaging. To induce shRNA-mediated depletion of Kif4A and simultaneous expression of its respective RNAi-resistant variants, 1 μg/ml of doxycycline (Sigma-Aldrich) was added to the refreshed medium each day for 3 days.

### Mitosis with Unreplicated genomes (MUGs)

MUGs was induced as described previously (Schlegel & Pardee, 1986, Wise & Brinkley, 1997). Briefly, U2OS cells stably expressing PA-GFP/mCherry-α-tubulin (gift from R. Medema, NKI, Amsterdam, the Netherlands) were first synchronized in mitosis with 2 μM nocodazole (Sigma-Aldrich) for 16 h. Mitotic cells were collected by shake-off, washed in phosphate-buffered saline (PBS) and plated at a density 4.5 × 10^5^ cells in 35 mm glass-bottomed dish (14mm, No. 1.5, MatTek Corporation). To inhibit DNA replication, 2 mM hydroxyurea (HU) (Sigma-Aldrich) was added directly to the cell culture medium for 23 h. Next day, the medium was replaced with fresh medium containing 2 mM HU and 5 mM caffeine and incubated for an additional 6 h.

### RNAi interference

U2OS cells stably expressing PA-GFP/mCherry-α-tubulin (van Heesbeen, Tanenbaum et al., 2014) were transfected with 50 nM siRNAs for 48-72 h, using Lipofectamine RNAiMAX (Thermo Fisher Scientific). The siRNAs used in this study are: KIF4A, 5′- GCAGAUUGAAAGCCUAGAG-3′ (Wandke et al., 2012); KIF2A, 5′- GGCAAAGAGAUUGACCUGG-3′ (Ganem & Compton, 2004); hKID, 5‘- CAAGCUCACUCGCCUAUUGTT-3′ (Wandke et al., 2012, Wolf, Wandke et al., 2006); KIF15, 5′-GGACAUAAAUUGCAAAUACTT-3′ (Tanenbaum et al., 2009); CLASP1, 5′- GGAUGAUUUACAAGACUGG-3′ (Mimori-Kiyosue, Grigoriev et al., 2006); CLASP2, 5′- GACAUACAUGGGUCUUAGA-3′ (Mimori-Kiyosue et al., 2006); NDC80, 5′- GAAUUGCAGCAGACUAUUA-3′; NuMA, 5′-AAGGGCGCAAACAGAGCACUA-3′ (Kotak, Busso et al., 2012); HSET, 5′-UCAGAAGCAGCCCUGUCAA-3′ (Cai et al., 2009); PRC1, 5′- AAAUAUGGGAGCUAAUUGGGA-3′ (Mollinari et al., 2002); MCAK, 5’- GAUCCAACGCAGUAAUGGU-3’ (Cassimeris & Morabito, 2004) and non-targeting control siRNA (D-001810-01-05, Dharmacon Inc.), 5′-UGGUUUACAUGUCGACUAA-3′. For rescue experiments of HSET, U2OS cells stably expressing mEOS-α-tubulin (Wandke et al., 2012) were transfected with 100 nM siRNA targeting 3′UTR for HSET, 5′- CAUGUCCCAGGGCUAUCAAAU-3′ (Cai et al., 2009).

### Immunoblotting and immunofluorescence

For immunoblotting, cells following subjected treatments were collected and lysed in NP40 buffer (50mM Tris-HCl pH 8, 150mM NaCl, 5mM EDTA, 0.5% NP-40, 1x EDTA-free protease inhibitor (Sigma-Aldrich), 1x phosphatase inhibitor cocktail (Sigma-Aldrich), 1mM PMSF) by two cycles of freezing and thawing using liquid N_2_. Protein extracts collected after centrifugation were subjected to SDS-PAGE and transferred onto nitrocellulose membrane (Bio-Rad). After blocking with PBS containing 0.1% Tween-20 and 5% milk, membranes were incubated overnight with primary antibodies (see Key Resources Table), washed in PBS containing 0.1% Tween-20 and then incubated 1 h with HRP-conjugated secondary antibodies (Jackson ImmunoResearch). After washing with PBS containing 0.1% Tween-20, immunodectection was performed with ECL (Bio-Rad). Immunoblots were performed using the following primary antibodies: rabbit anti-KIF15 (1:1000; Cytoskeleton Inc., Cat#AKIN13), rabbit anti-KIF2A (1:1000; Novus Biologicals, NB500-180); rabbit anti-KIF4A (1:1000) (Wandke et al., 2012), mouse anti-hKID (8C12) (1:1000) (Wandke & Geley, 2006); rat anti-CLASP1 (1:200) and rat anti-CLASP2 (1:200) (Maffini et al., 2009), mouse anti-PRC1 (C-1) (1:500; Santa Cruz, sc-3769839), mouse anti-NDC80/HEC1 (1:2000; Nordic Biosite, GTX70268), mouse anti-NuMA (F-11) (1:250; Santa Cruz, sc-365532), rabbit anti-HSET (1:500; Bethyl, A300-951A), mouse anti-MCAK (1:1000; Abnova H00011004-M01), mouse anti-Vinculin (Vin-11-5) (1:5000; Sigma-Aldrich, SAB4200729, Vin-11-5), mouse anti-α-tubulin (B-5-1-2) (1:10000; Sigma-Aldrich T5168, B-5-1-2), mouse anti-GAPDH (1:25000; Proteintech 60004-1-Ig), HRP-conjugated secondary antibodies (1:10000; Jackson ImmunoResearch), and visualized by ECL (Bio-Rad). For immunofluorescence, U2OS cells grown on glass coverslips were fixed either by ice-cold methanol for 4 min at −20°C, or by 4% paraformaldehyde in PHEM buffer for 20 min at 37°C as described before (DeLuca, 2010). Cells were washed with 1x phosphate-buffered saline (PBS) and immunostained with primary and Alexa Fluor-conjugated secondary antibodies (see Key Resources Table) diluted in IF stain (1x PBS, 1% FBS, 0.5% Tween). DNA was counterstained with DAPI (Sigma-Aldrich, final concentration 0.1 μg/ml) and mounted on glass slides using Fluoromount-G mounting media (Southern Biotech). Images were acquired using LSM700 confocal microscope (Carl Zeiss Microimaging Inc.) mounted on a Zeiss-Axio imager Z1 equipped with plan-apochromat 63 × /1.40 oil DIC M27 objective (Carl Zeiss, Inc.) and Zen 2008 software (Carl Zeiss, Inc.). Primary antibodies used were: mouse anti-α-tubulin (DM1A) (1:2000; Sigma-Aldrich, T9026), rabbit anti-α-tubulin (1:500; Abcam, ab15246); rabbit anti-KIF4A (1:500; Bethyl, A301-074A), rabbit anti-CENP-E (1:500, Abcam, ab133583), mouse anti-PRC1 (1:250; Santa Cruz, sc-3769839). The Alexa Fluor-conjugated secondary antibodies (1:1000; Thermo Fisher Scientific) were used for secondary antibody labeling and DAPI (1 μg/mL) as DNA counterstain.

### Live-cell imaging

Time-lapse imaging was performed in a heated incubation chamber at 37°C with controlled humidity and 5% CO_2_ supply using a Plan-Apochromat 63x/1.4NA oil objective with differential interference contrast mounted on an inverted Zeiss Axio Observer Z1 microscope (Marianas Imaging Workstation (3i - Intelligent Imaging Innovations, Inc., Denver, CO, USA), equipped with a CSU-X1 spinning-disk confocal head (Yokogawa Corporation of America) and four laser lines (405 nm, 488 nm, 561nm and 640 nm). Images were acquired using an iXon Ultra 888 EM-CCD camera (Andor Technology). Fifteen 1μm-separated z-planes were collected every 1 min for 2 h. To quantify the spindle length, the distance between mCherry-labeled spindle poles was measured in 3D using ImageJ (National Institute of Health, Bethesda, MD, USA). For MCAK RNAi-mediated recovery experiments, the spindle length was quantified using time-lapse images of mitotic spindles with fifteen 1μm-separated z-planes imaged every 2min.

### Photoactivation and MT-flux analysis

For the MT poleward flux measurement, photoactivation and photoconversion experiments were performed using U2OS-PA-GFP/mCherry-α-tubulin, U2OS-PA-GFP and U2OS-mEOS-α-tubulin cell lines, respectively, cultured in 35 mm glass-bottomed dishes (14mm, No. 1.5, MatTek Corporation). Fluorescent signals of SiR-DNA and mCherry-α-tubulin were used to differentiate between early prometaphase and metaphase cells. Two transversal line-shaped regions of interest, placed perpendicular to the main spindle axis on both sides of the metaphase plate, were photoactivated in bipolar spindles, while 2-3 concentric rings were used for photoactivation of monopolar spindles. Photoactivation was carried out by one 5 ms pulse from a 405 nm laser. Images were acquired every 5 s for 4 min over three 0.5 μm-separated z-planes. Kymograph generation and velocity quantification are performed using a custom-written Matlab script (Pereira & Maiato, 2010). Initially, the spindle is stabilized by tracking the spindle poles, the coordinates of which are looked up to rotate and translate the ROIs for the definition of each kymograph layer. Each layer (a matrix) is then collapsed into a vector using a projection criteria (maximum or sum), generating a collapsed kymograph such as the ones shown in Fig. 1d. To calculate flux velocity, a sparse set of manually chosen points is used as a first guess for stripe position at specific points. This initial estimate is made continuous by computing a cubic spline curve. Within a preset neighborhood, the position of the stripe at each time point is then corrected by calculating an intensity-based centroid. Finally, a linear fit is performed to these corrected positions, the slope of which is the stripe velocity averaged over the time window used to generate the kymograph. In fact, two stripes are imprinted in each cell, the velocities of which (one positive and one negative) are subtracted and then halved to yield a poleward flux velocity relative to a virtual spindle equator, which is used as the final readout presented in quantifications.

### CH-STED microscopy

U2OS cells grown on glass coverslips were fixed using a solution of 4% paraformaldehyde (Electron microscopy sciences) and 0.1% glutaraldehyde (Electron microscopy sciences) for 10 min at RT. Autofluorescence was quenched using freshly prepared 0.01% sodium borohydride for 7 min at RT. Cells were permeabilized with 0.5% Triton-X100 in PBS for 10 min at RT, washed, and incubated in blocking solution (10% fetal bovine serum (FBS) in 0.05% PBS-Tween) for 30 min. Immunolabelling was performed with the following primary antibodies: rabbit anti-CENP-E (1:25), rabbit anti-KIF4A (1:50; PA5-30492), rat anti-tyrosinated tubulin (1:100) and guinea pig anti-CENP-C (1:500), fluorescently labelled secondary antibodies anti-rabbit 580 STED (1:100; Abberior), anti-rat STAR RED (1:100, Abberior) and anti-guinea pig Alexa Fluor-488 (1:1000, Thermo Fisher Scientific) and DAPI (1 μg/mL) as DNA counterstain.CH-STED microscopy was performed as described previously (Pereira et al., 2019). An Abberior Instruments ‘Expert Line’ gated-STED was used coupled to a Nikon Ti microscope with 60x 1.4NA Plan-Apo objective (Nikon, Lambda Series) oil-immersion, pinhole size of 0.8 Airy units, 40 MHz modulated excitation (405, 488, 560 and 640nm) and depletion (775nm) lasers. Abberior’s Inspector software was used to control the acquisition process. CH-STED was always acquired after 2D-STED (and z-STED) acquisitions to avoid artifacts caused by photo-bleaching.

### Quantification and Statistical Analysis

Statistical analysis and graphs were generated in GraphPad Prism. Statistical details of experiments and tests used are indicated in the figure legends. Statistical significance was determined by the Student’s t-test or one-way ANOVA where the conditions were compared with control. Details of the statistical significance and n values for each conditions can be found in the figures and figure legends.

## Acknowledgments

We thank Duane Compton and Rene Medema for providing the U2OS PA-GFP-tubulin and U2OS PA-GFP-tubulin/mCherry-tubulin cell lines, respectively. We thank Claire Walczak for providing the GFP-HSET and GFP-HSET N593K plasmids. We thank Martina Barisic for technical assistance. Work in the lab of MB is supported by grants from Danish Cancer Society Scientific Committee (KBVU; R146-A9322) and the Lundbeck Foundation (R215-2015-4081). Work in the lab of HM is funded by the European Research Council (grant agreement No 681443) under the European Union’s Horizon 2020 research and innovation program.

## Author contributions

YS, GR, AJP, HM and MB designed experiments; YS generated tools and performed and analyzed most of the experiments; YS and MB performed photoactivation experiments, image quantification and analysis. GR performed and analyzed the spindle size-related experiments. MO performed and analyzed initial photoactivation experiments; YS, AJP and HM performed CH-STED experiments and analysis; SG provided reagents; SE contributed to designing and analyzing the experiments; MB, YS and GR wrote the manuscript, with contributions from all authors; MB conceived and coordinated the project.

## Expanded View Figure Legends

**Figure EV1. Efficient RNAi-mediated knockdown of selected MT-flux driving candidates and the effect of KIF4A overexpression on MT-flux. A.** Immunoblot analysis of lysates from U2OS cells stably expressing PA-GFP/mCherry-α-tubulin treated with control or indicated siRNAs. Antibodies against respective proteins were used to validate siRNA mediated knockdown with GAPDH, α-tubulin and vinculin serving as the loading controls. **B.** Immunoblot analysis of cell lysates obtained from U2OS cells stably expressing PA-GFP-α-tubulin infected with different multiplicity of infection (MOI) ratios of mCherry-KIF4A expressing adenovirus. Expression levels of KIF4A upon adenoviral titration was detected using anti-KIF4A antibody, with GAPDH as loading control. **C.** Quantification of the MT-flux rates upon KIF4A overexpression. The individual values and their mean ± SD are plotted. N (number of cells, number of independent experiments): Control (23, 3); 1 MOI (33, 3); 2 MOI (26, 3), 4 MOI (9, 1). P-values were calculated using one-way ANOVA. * P ≤ 0.05, *** P ≤ 0.001. **D.** Schematic model illustrating KIF4A-mediated forces in MT-flux driving. Forces exerted by KIF4A motors promote motion of chromosomes relative to MTs. Once a bi-oriented chromosome aligns at the metaphase plate, KIF4A forces can no longer do work on it. Rather, the reactive force on MTs may promote non-KT-MTs to flux towards the poles.

**Figure EV2. Analysis of the roles of MT-crosslinking proteins NuMA, HSET and PRC1 in MT-flux. A.** Immunoblot analysis of the respective knockdown efficiencies in U2OS cells stably expressing PA-GFP/mCherry-α-tubulin treated with control or indicated siRNAs. GAPDH, α-tubulin and vinculin were used as a loading control. **B.** Spinning disk confocal time series images of photoconverted mEOS-α-tubulin (red) in U2OS cells stably expressing mEOS-α-tubulin (cyan). White arrowheads highlight the photoconverted regions fluxing towards the poles. Scale bar, 10 μm. Time, min:sec. **C.** Immunoblot analysis of U2OS cells stably expressing mEOS-α-tubulin transfected with control or HSET 3′UTR-targeting siRNAs and co-transfected with the RNAi-resistant GFP-HSET constructs. Expression levels of HSET was observed using anti-HSET antibody with α-tubulin serving as a loading control.

**Figure EV3. Analysis of CENP-E contribution to MT-flux in the absence of stable end-on KT-MT attachments. A.** Immunoblot analyses of the efficiency of knockdown from U2OS cells stably expressing PA-GFP/mCherry-α-tubulin treated with control or indicated siRNAs, with GAPDH or α-tubulin used as a loading control. **B.** Representative point-scanning confocal maximum intensity projections of control and MUGs mitotic spindles in U2OS cells, immunostained with antibodies against CENP-E and α-tubulin. DNA was counterstained with DAPI. CENP-E in red, α-tubulin in green and DNA in blue in merged image. Scale bar, 10 μm.

**Figure EV4. Distribution of antiparallel interpolar MTs during mitosis. A.** Representative spinning disk confocal time series of MT-flux during prometaphase (upper panel) and metaphase (lower panel) in U2OS cells stably co-expressing PA-GFP-α-tubulin (cyan) and mCherry-α-tubulin (red), with chromosomes labelled using SiR-DNA (grey). White arrowheads highlight that the photoactivated regions in prometaphase split and move towards both poles (upper panel), whereas metaphase spindles flux uniformly towards individual poles (lower panel). Scale bars, 10 μm. Time, min:sec. **B.** Representative maximum projections of point-scanning confocal immunofluorescence images of prometaphase and metaphase spindles in U2OS cells, stained using PRC1 and α-tubulin antibodies. DNA was counterstained with DAPI. PRC1 (cyan) and α-tubulin (red) are shown in merged image. The white line signifies area of PRC1-stained antiparallel MTs within the spindle. Scale bar, 10 μm.

**Figure EV5. MT flux counterbalances MCAK activity to maintain spindle length. A.** Illustration of the relationship between MT-flux and spindle length in the presence or absence of stable KT-MT end-on attachments. **B.** Immunoblot analysis of the knockdown efficiency of indicated proteins in U2OS PA-GFP/mCherry-α-tubulin cells treated with control or target specific siRNAs. GAPDH and α-tubulin serve as a loading controls. **C.** Percentage of cells where bipolar spindle collapses to become monopolar after indicated treatments. N (number of cells, number of independent experiments): siKIF15 + STLC + siKIF4A (59, 3), siKIF15 + STLC + siKIF4A +siMCAK (94, 3). P-values were calculated using Student’s t test. * P ≤ 0.05.

**Table EV1.** MT-flux and mitotic spindle length of cells following depletion or inhibition of indicated targets. The flux rate and spindle length values are mean ± s.d.

| **Bipolar mitotic spindles** | **Average metaphase flux rate ($\mu\text{m}/\text{min}$)** | **Average spindle length ($\mu\text{m}$)** | **No. of cells (No. of experiments)** |
| --- | --- | --- | --- |
| siControl | $0.61 \pm 0.22$ | $13.30 \pm 1.79$ | 49 (5) |
| 5$\mu\text{M}$ STLC | $0.58 \pm 0.12$ | $14.89 \pm 1.27$ | 31 (3) |
| siKIF15/kinesin-12 | $0.6 \pm 0.15$ | $11.66 \pm 0.94$ | 39 (3) |
| siKIF15 + STLC | $0.43 \pm 0.11$ | $11.76 \pm 1.56$ | 29 (3) |
| siKID/kinesin-10 | $0.59 \pm 0.2$ | $14.04 \pm 1.35$ | 32 (3) |
| siKIF4A/kinesin-4 | $0.32 \pm 0.14$ | $11.41 \pm 1.02$ | 28 (3) |
| siKIF15 + siKIF4A + STLC | $0.15 \pm 0.10$ | $8.38 \pm 1.58$ | 10 (3) |
| shKIF4A (uninduced) | $0.59 \pm 0.10$ | | 48 (4) |
| shKIF4A | $0.29 \pm 0.13$ | | 60 (4) |
| shKIF4A + KIF4A WT (uninduced) | $0.57 \pm 0.10$ | | 36 (5) |
| shKIF4A + KIF4A WT | $0.53 \pm 0.15$ | | 43 (5) |
| shKIF4A + KIF4A K94A (uninduced) | $0.63 \pm 0.15$ | | 30 (4) |
| shKIF4A + KIF4A K94A | $0.33 \pm 0.14$ | | 32 (4) |
| shKIF4A + KIF4A delZip1 (uninduced) | $0.63 \pm 0.13$ | | 32 (4) |
| shKIF4A + KIF4A delZip1 | $0.30 \pm 0.10$ | | 30 (4) |
| Ctrl OX | $0.56 \pm 0.09$ | | 23 (3) |
| KIF4A – 1 MOI | $0.69 \pm 0.14$ | | 33 (3) |
| KIF4A – 2 MOI | $0.65 \pm 0.14$ | | 26 (3) |
| KIF4A – 4 MOI | $0.74 \pm 0.18$ | | 9 (1) |
| siKIF2A/kinesin-13 | $0.17 \pm 0.12$ | $11.03 \pm 1.55$ | 36 (3) |
| siCLASPs | $0.27 \pm 0.18$ | $10.15 \pm 1.5$ | 21 (2) |
| 200nM GSK923295 | $0.52 \pm 0.21$ | $14.39 \pm 1.41$ | 44 (3) |
| siPRC1 | $0.61 \pm 0.22$ | $15.03 \pm 2.37$ | 45 (3) |
| siNuMA | $0.48 \pm 0.13$ | $15.34 \pm 1.38$ | 40 (3) |
| siHSET/kinesin-14 | $0.36 \pm 0.13$ | $14.82 \pm 1.53$ | 37 (3) |
| siNuMA + siHSET | $0.35 \pm 0.16$ | $14.87 \pm 1.46$ | 39 (3) |
| siControl (mEOS) | $0.61 \pm 0.13$ | | 23 (3) |
| siHSET (3'UTR) | $0.37 \pm 0.13$ | | 33 (3) |
| siHSET (3'UTR) + HSET WT | $0.6 \pm 0.08$ | | 29 (5) |
| siHSET (3'UTR) N593K | $0.55 \pm 0.11$ | | 30 (5) |
| siNDC80 | $0.91 \pm 0.2$ | $15.97 \pm 3.10$ | 59 (6) |
| siNDC80 + siKIF4A | $0.89 \pm 0.24$ | $16.74 \pm 2.19$ | 41 (3) |
| siNDC80 + STLC | $0.92 \pm 0.18$ | $16.64 \pm 2.76$ | 28 (3) |
| siNDC80 + siCLASPs | $0.86 \pm 0.21$ | $17.66 \pm 2.17$ | 33 (3) |
| siNDC80 + siKIF2A | $0.64 \pm 0.22$ | $15.78 \pm 3.15$ | 38 (3) |
| siNDC80 + siKIF15 | $0.97 \pm 0.18$ | $18.06 \pm 2.35$ | 36 (4) |
| siNDC80 + GSK923295 | $0.6 \pm 0.28$ | $15.21 \pm 1.96$ | 38 (3) |
| siNDC80 + siKIF15 + STLC | $0.52 \pm 0.13$ | $16.88 \pm 2.90$ | 33 (5) |
| siNDC80 + siKIF15 + STLC + GSK923295 | $0.10 \pm 0.09$ | $13.83 \pm 3.63$ | 25 (3) |
| Early prometaphase | $0.84 \pm 0.15$ | $13.47 \pm 1.99$ | 23 (3) |
| Early prometaphase + GSK923295 | $0.54 \pm 0.18$ | $13.40 \pm 1.86$ | 28 (3) |
| Early prometaphase + GSK923295 + siKIF15 + STLC | $0.07 \pm 0.11$ | $8.37 \pm 1.80$ | 11 (2) |
| Ctrl (MUGs) | $0.5 \pm 0.13$ | | 30 (3) |
| MUGs | $0.31 \pm 0.13$ | | 34 (3) |
| **Monopolar mitotic spindles** | **Average flux rate ($\mu\text{m}/\text{min}$)** |  | **No. of cells (No. of experiments)** |
| siControl | $0.50 \pm 0.14$ | | 35 (3) |
| siKIF15 | $0.48 \pm 0.11$ | | 44 (3) |
| siKID | $0.44 \pm 0.11$ | | 38 (3) |
| siKIF4A | $0.33 \pm 0.11$ | | 44 (3) |
| siKIF2A | $0.17 \pm 0.09$ | | 38 (3) |
| siCLASPs | $0.28 \pm 0.07$ | | 12 (3) |
| GSK923295 | $0.34 \pm 0.07$ | | 33 (3) |
| siNDC80 | $0.48 \pm 0.08$ | | 47 (3) |

## Expanded view movie legends

**Movie EV1. MT-flux in KIF4A-depleted cells.** U2OS cells stably expressing PA-GFP-α-tubulin (cyan) and mCherry-α-tubulin (red) were transfected with control siRNAs (left) or KIF4A siRNAs (right). 405nm laser-based photoactivation was performed to monitor MT-flux using spinning disk confocal microscopy. Time, min:sec.

**Movie EV2. MT-flux in STLC-induced monopolar spindles.** U2OS cells stably expressing PA-GFP-α-tubulin (cyan) and mCherry-α-tubulin (red) were treated with STLC alone (left) or in combination with taxol (right). 405nm laser-based photoactivation was performed to monitor MT-flux using spinning disk confocal microscopy. Time, min:sec.

**Movie EV3. Mitosis in cells expressing mCherry-KIF4A WT, mCherry-KIF4A K64A motor mutant and mCherry-KIF4A ΔZip1.** U2OS cells stably expressing PA-GFP-α-tubulin were induced to co-express KIF4A shRNA and RNAi-resistant mCherry-KIF4A variants (right). Chromosomes were stained using SiR-DNA (left). Live-cell imaging was performed using spinning disk confocal microscopy. Time, hours:min.

**Movie EV4. MT-flux in cells undergoing mitosis with unreplicated genomes (MUGs).** U2OS cells stably expressing PA-GFP-α-tubulin (cyan) and mCherry-α-tubulin (red) were treated with caffeine only as control (left) or with hydroxyurea and caffeine to induce MUGs (right). 405nm laser-based photoactivation was performed to monitor MT-flux using spinning disk confocal microscopy. Chromosomes were stained using SiR-DNA (grey). Time, min:sec.

**Movie EV5. MT-flux in the absence of MT-crosslinking proteins HSET and NuMA.** U2OS cells stably expressing PA-GFP-α-tubulin (cyan) and mCherry-α-tubulin (red) were transfected with control siRNAs (left) and siRNAs against HSET and NuMA (right). 405nm laser-based photoactivation was performed to monitor MT-flux using spinning disk confocal microscopy. Time, min:sec.

**Movie EV6. The role of CENP-E in MT-flux in the absence of stable end-on KT-MT attachments.** U2OS cells stably expressing PA-GFP-α-tubulin (cyan) and mCherry-α-tubulin (red) were transfected with control siRNAs (left), NDC80 siRNAs (middle) and NDC80 siRNAs co-treated with CENP-E inhibitor GSK923295 (right). 405nm laser-based photoactivation was performed to monitor MT-flux using spinning disk confocal microscopy. Time, min:sec.

**Movie EV7. MT-flux in prometaphase vs. metaphase cells.** Live prometaphase (upper panel) and metaphase (lower panel) U2OS cells stably expressing PA-GFP-α-tubulin (cyan) and mCherry-α-tubulin (red) were imaged using spinning disk confocal microscopy. Chromosomes were stained using SiR-DNA (left, grey). 405nm laser-based photoactivation was performed to monitor MT-flux. Time, min:sec.

**Movie EV8. MT-flux in cells with simultaneous inactivation of three motors.** U2OS cells stably expressing PA-GFP-α-tubulin (cyan) and mCherry-α-tubulin (red) were transfected with either control siRNAs or siRNAs against NDC80 (left) and siRNAs against indicated motors (right). 405nm laser-based photoactivation was performed to monitor MT-flux using spinning disk confocal microscopy. Time, min:sec.

