## Supplementary figures and images for "Microtubule poleward flux in human cells is driven by the coordinated action of four kinesins"

### Supplemental data

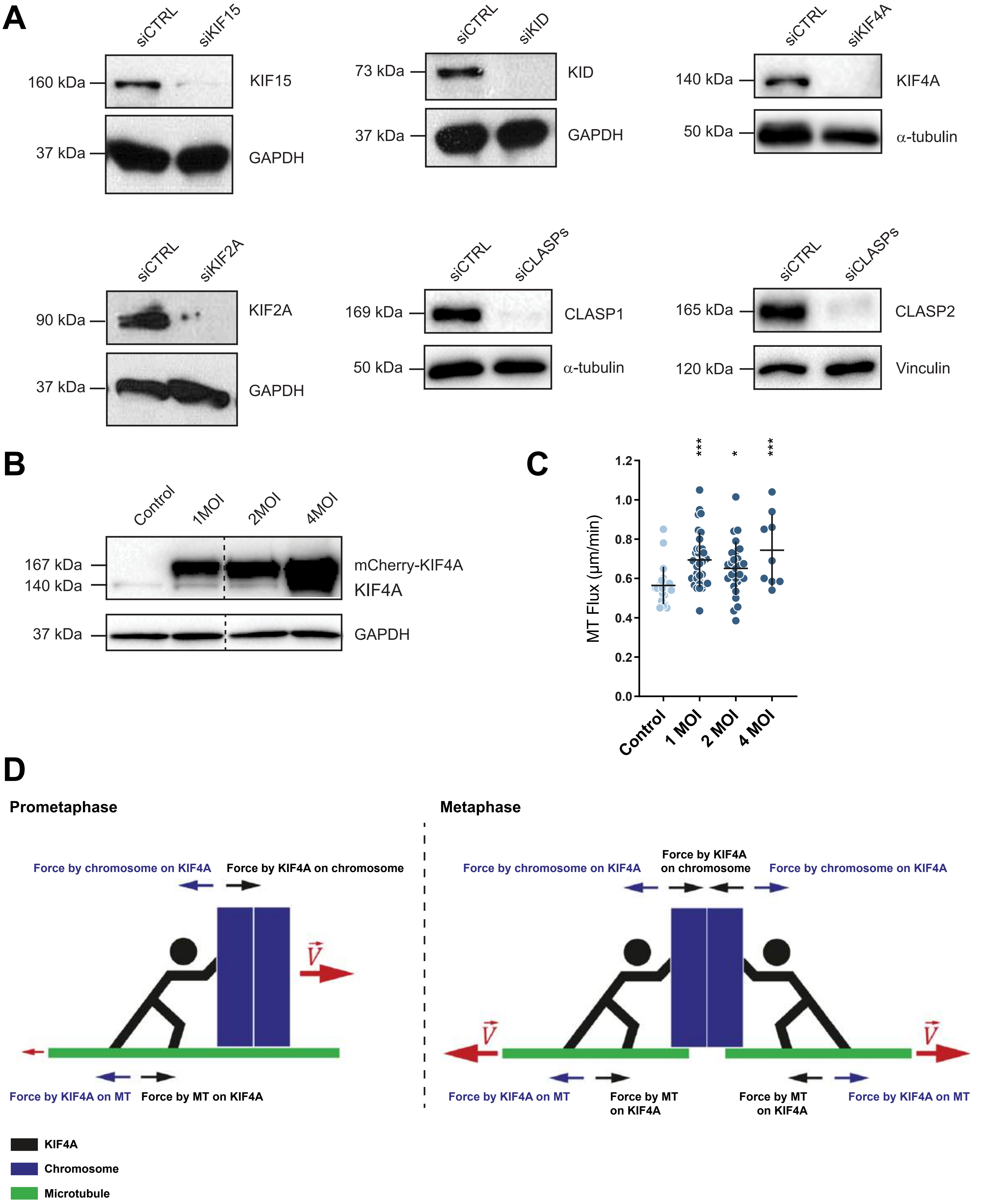

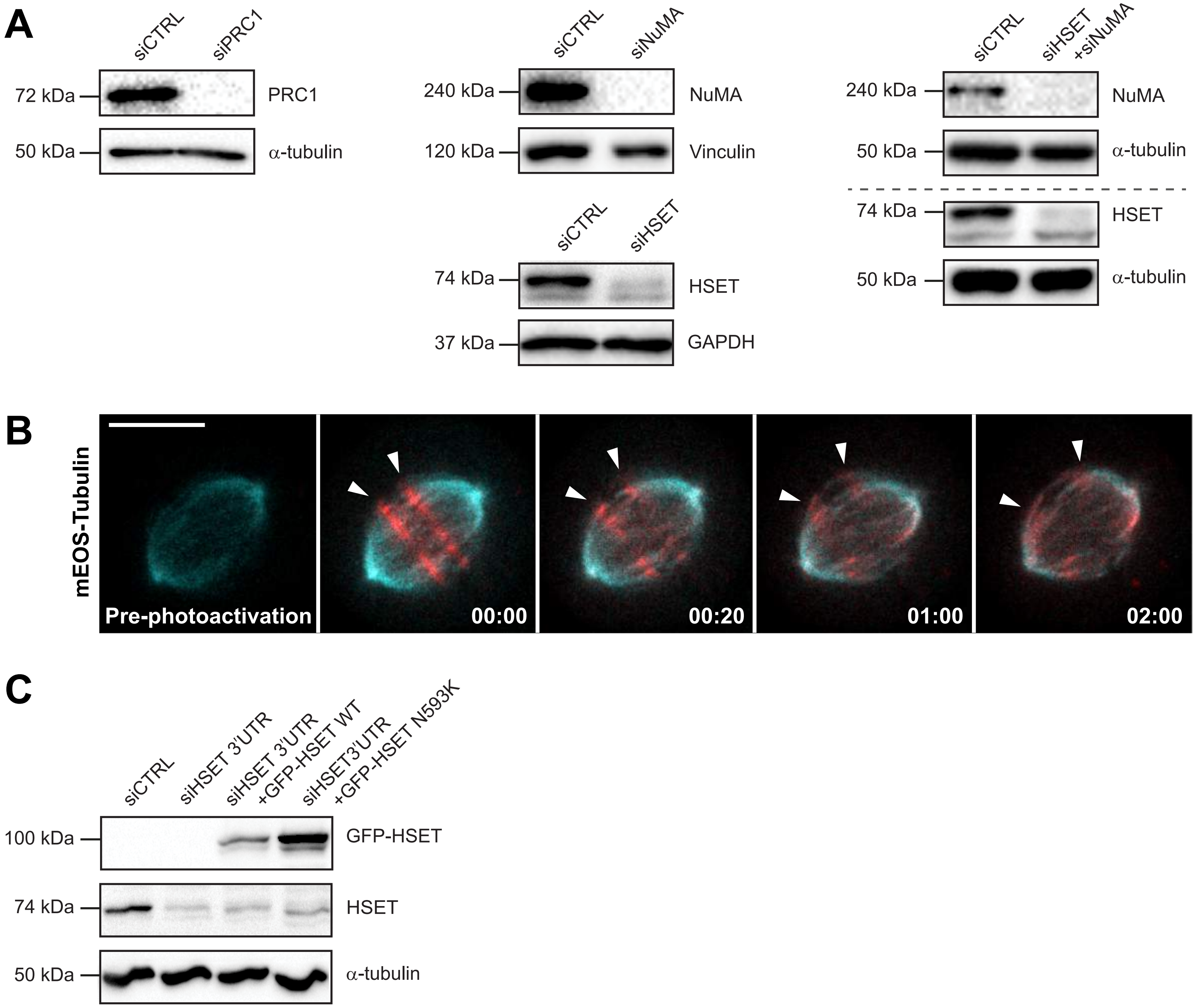

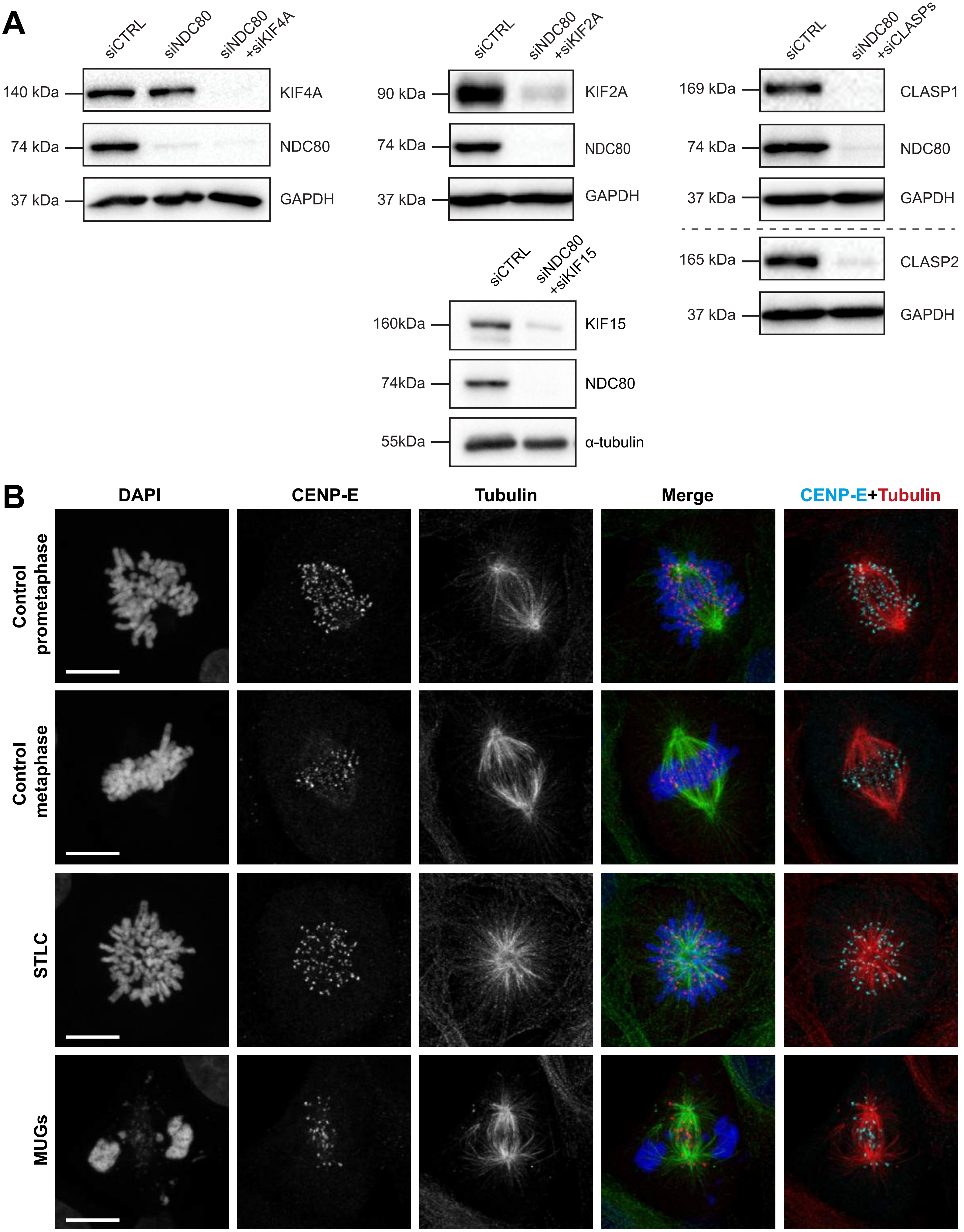

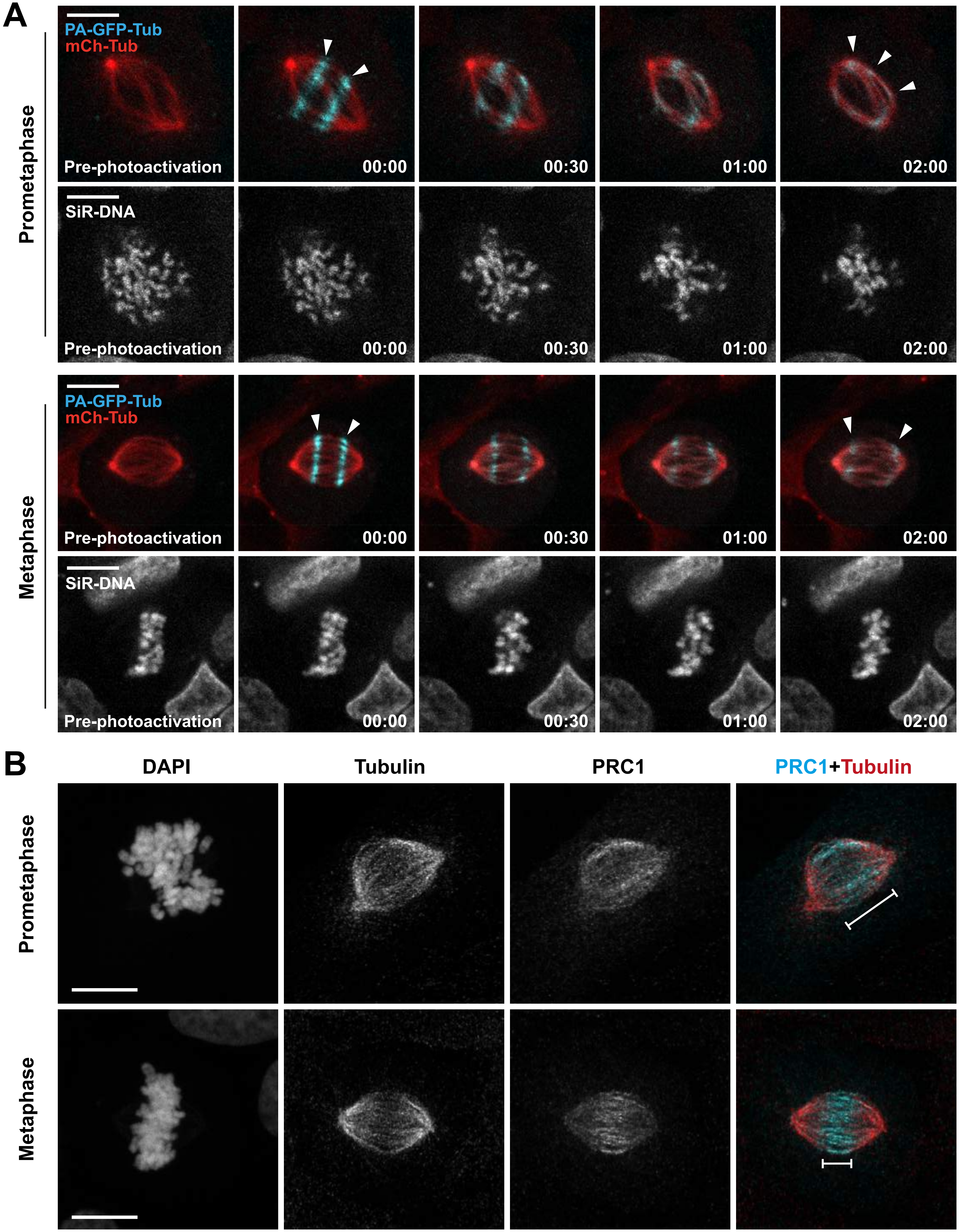

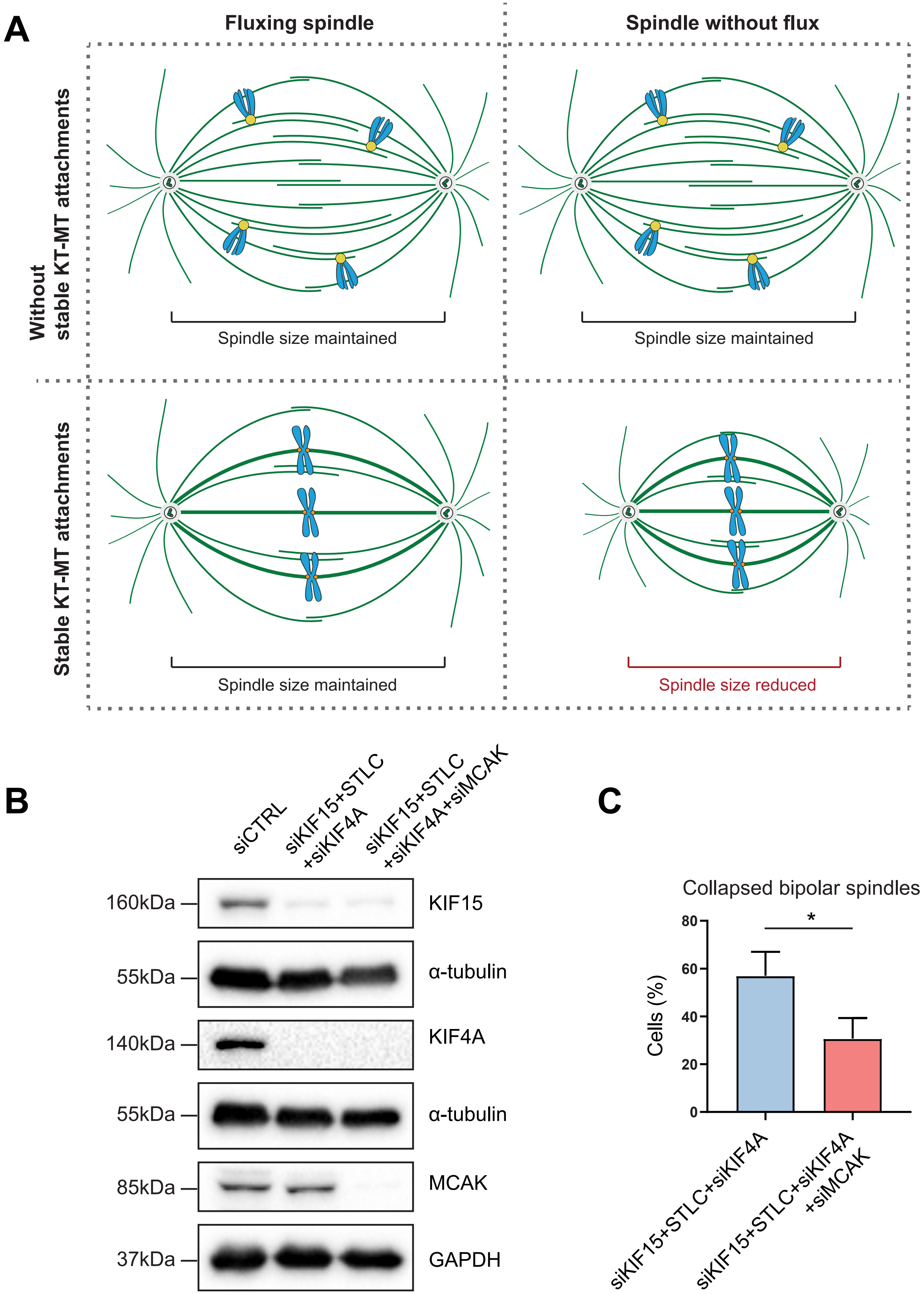
